# Molecular insights into ago-allosteric modulation of the human glucagon-like peptide-1 receptor

**DOI:** 10.1101/2021.05.29.446269

**Authors:** Zhaotong Cong, Li-Nan Chen, Honglei Ma, Qingtong Zhou, Xinyu Zou, Chenyu Ye, Antao Dai, Qing Liu, Wei Huang, Xianqiang Sun, Xi Wang, Peiyu Xu, Lihua Zhao, Tian Xia, Wenge Zhong, Dehua Yang, H. Eric Xu, Yan Zhang, Ming-Wei Wang

**Author notes:** These authors contributed equally: Zhaotong Cong, Li-Nan Chen, Honglei Ma and Qingtong Zhou. Correspondence: Dehua Yang or H. Eric Xu or Yan Zhang or Ming-Wei Wang.

## Abstract

The glucagon-like peptide-1 (GLP-1) receptor is a validated drug target for metabolic disorders. Ago-allosteric modulators are capable of acting both as agonists on their own and as efficacy enhancers of orthosteric ligands. However, the molecular details of ago-allosterism remain elusive. Here, we report three cryo-electron microscopy structures of GLP-1R bound to (i) compound 2 (an ago-allosteric modulator); (ii) compound 2 and GLP-1; and (iii) compound 2 and LY3502970 (a small molecule agonist), all in complex with heterotrimeric G_s_. The structures reveal that compound 2 is covalently bonded to C347 at the cytoplasmic end of TM6 and triggers its outward movement in cooperation with the ECD whose N terminus penetrates into the GLP-1 binding site. This allows compound 2 to execute positive allosteric modulation through enhancement of both agonist binding and G protein coupling. Our findings offer the structural basis of ago-allosterism at GLP-1R and new knowledge to design better therapeutics.

## Introduction

As one of the most well-known class B1 G-protein-coupled receptors (GPCRs), the glucagon like peptide-1 receptor (GLP-1R) is a clinically validated drug target for type 2 diabetes and obesity^1–3^. Despite the therapeutic success, currently available GLP-1R agonists are suboptimal due to poor patient compliance caused by subcutaneous injections and several side-effects such as nausea and vomiting^4,5^. Oral-delivery of GLP-1 mimetics have been pursued for many years, with semaglutide being the first and only one in the clinic albeit its low bioavailability and frequent gastrointestinal complaints^6,7^. Thus, development of orally active small molecule GLP-1R modulators remains an attractive strategy and a few compounds have progressed to clinical trials^8^.

The scientific advances in GPCR structural biology, fueled by tremendous interests from both academia and industry, led to 16 solved GLP-1R structures. In 2017, two inactive structures bound to negative allosteric modulators (NAMs) ^9^, one intermediate structure bound to a truncated peptide agonist^10^ and one active structure in complex with GLP-1 and heterotrimeric G_s_^11^ were reported. This was followed by eight G protein-bound structures in complex with peptides (GLP-1 and exendin-P5)^12,13^, non-peptidic agonists (TT-OAD2, PF-06882961, CHU-128, RGT1383 and LY3502970)^13–16^ and a positive allosteric modulator (PAM) LSN3160440 (ref. ^17^), as well as a peptide-free *apo* state^18^ and three thermal-stabilized TMD^9^ structures. They display diversified conformations and reveal key information on ligand recognition and GLP-1R activation. Integrated with the functional, pharmacological and computational data, these studies demonstrate distinct locations and different conformations of the orthosteric binding pocket between small molecule and peptidic ligands.

Allosteric modulation by PAMs that bind anywhere distinct from the orthosteric site of endogenous ligands and enhance the activity of agonist in an uncompetitive way is an alternative approach to peptide therapy^19–22^. Previous studies showed that several PAMs of GLP-1R, such as the most characterized electrophilic chemotypes-compound BETP^23,24^ and the substituted quinoxalines compound 2 (6,7-dichloro-3-methanesulfonyl-2-tert-butylamino-quinoxaline)^25,26^, were able to initiate or promote receptor activation^9,27^. Apart from increasing the binding affinity of GLP-1, compound 2 also acts as an agonist on its own, thereby being classified as an ago-allosteric modulator (ago-PAM) of GLP-1R^25,28^. Specifically, in the absence of an orthosteric ligand, compound 2 displayed a strong partial agonism in cAMP response (80% Emax of GLP-1), a weak partial agonism in pERK1/2 and no detectable calcium mobilization response^29,30^. It also caused less GLP-1R internalization than GLP-1, due to the weaker β-arrestin recruitment (30-50% Emax of GLP-1)^30^. As a PAM, compound 2 caused a concentration-dependent increase in the affinity of GLP-1 and oxydomodulin^31^, and induced biased signaling in a probe-dependent manner^31,32^. In addition, compound 2 was also reported to modulate cAMP responses elicited by small molecular agonists including Boc5, BMS21 and TT15 (ref. ^29^). Although structurally and functionally unique, this compound has poor pharmacokinetic properties due to its electrophilic nature that preclude potential clinical development^33^. Interestingly, like BETP, compound 2 interacts with cysteine 347^6.36^ via covalent modification at the cytoplasmic side of GLP-1R, consistent with our previous observation that an intact ECD is required for the activity of BETP and compound 2 (ref. ^34^) albeit the basis for such a requirement remains unknown. This distinct feature prompts us to use compound 2 as a tool to study the structural basis of its ago-allosterism in the context of both peptidic and non-peptidic ligands.

Here we report high-resolution cryo-electron microscopy (cryo-EM) structures of human GLP-1R–G_s_ in complex with compound 2 alone, with both compound 2 and endogenous peptidic agonist GLP-1, or with compound 2 and a small molecule agonist (LY3502970). The structural information obtained from this study provides valuable insights into molecular mechanisms by which compound 2 exerts the ago-allosteric action and its relevance to drug discovery.

## Results

### Structure determination

To understand activation and allosterism of compound 2 on GLP-1R, we determined the structures of GLP-1R–G_s_ complexes bound to compound 2 alone and in the presence of GLP-1 or LY3502970 (Fig. 1). The NanoBiT tethering strategy and Nb35 were used to stabilize the protein complexes^35,36^(Supplementary Fig. 1a). These protein complexes were then purified, resolved as monodispersed peaks on size-exclusion chromatography (SEC), and verified by SDS gel and negative staining to ascertain all the expected components (Supplementary Fig. 1c-e). Vitrified complexes were imaged by cryo-EM. After sorting by constitutive 2D and 3D classifications, 3D consensus density maps were reconstructed with global resolutions of 2.5Å (without ECD) or 3.3Å (with ECD) for compound 2 alone, 2.5Å for compound 2 plus GLP-1, and 2.9Å for compound 2 plus LY3502970, respectively (Supplementary Fig. 2 and Supplementary Table 1). The cryo-EM maps allowed us to build an unambiguous model for most regions of the complexes except for the flexible α-helical domain (AHD) of Gα and the stalk between transmembrane helice 1 (TM1) and extracellular domain (ECD), which were poorly resolved in most cryo-EM structures of GPCR–G_s_ complex (Supplementary Fig. 3). In addition, residues N338 to T343 in the intracellular loop 3 (ICL3) of the compound 2–GLP-1–GLP-1R–G_s_ and compound 2–LY3502970–GLP-1R–G_s_ complexes, F369 to R376 in the extracellular loop 3 (ECL3) of the compound 2–GLP-1R–G_s_ and compound 2–LY3502970–GLP-1R–G_s_ complexes, and P56 to F61 in the ECD of the compound 2–GLP-1R–G_s_ complex were also poorly resolved and thus omitted from the corresponding final model. The ECD in the GLP-1 and compound 2-bound complexes was clear enough to enable modeling of the backbone and a majority of side-chains using the ECD and TMD-refined maps (Fig. 1).

**Fig. 1.**
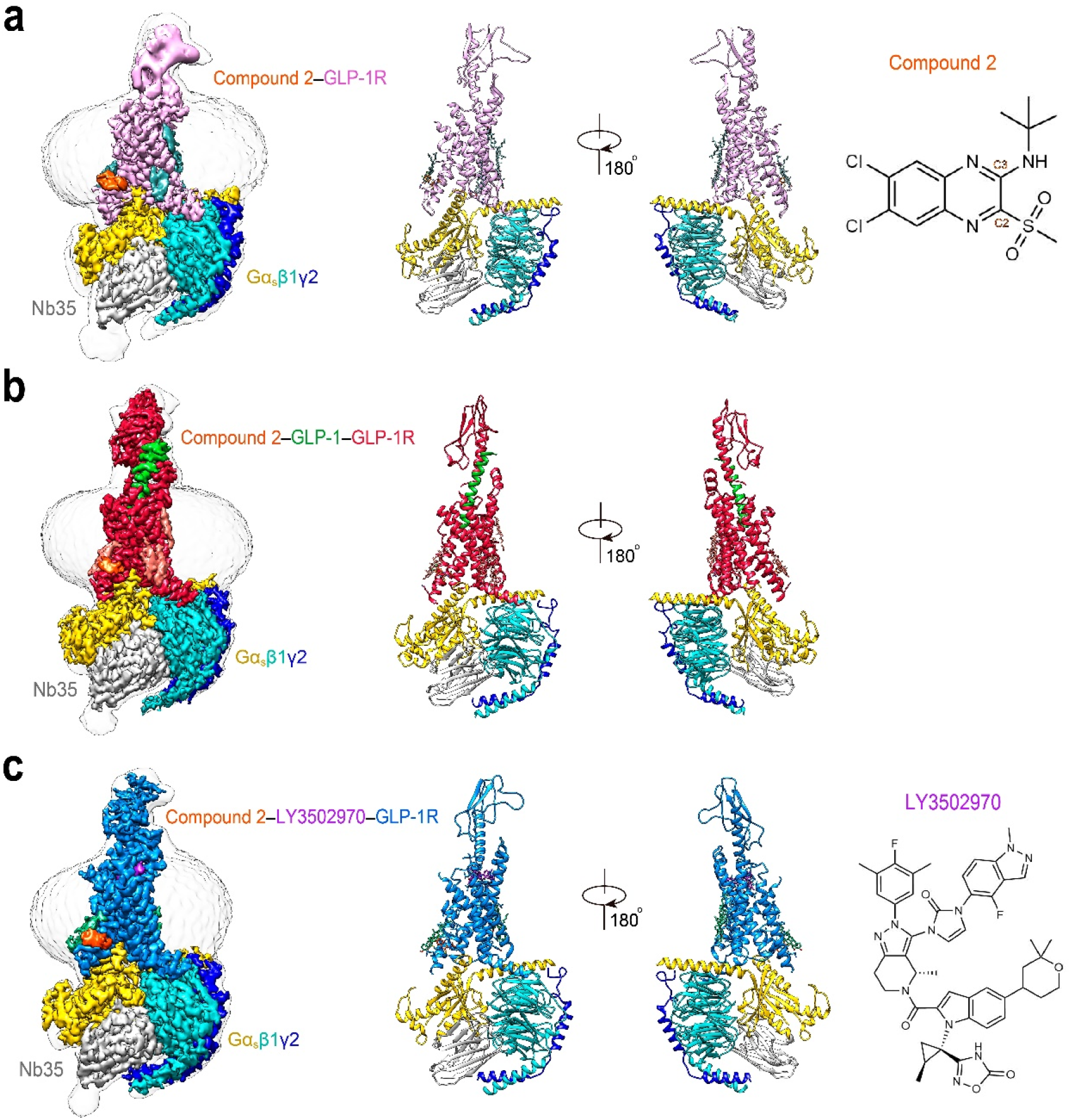
The overall cryo-EM structure of GLP-1R–G_s_ complexes. The cryo-EM maps with a discshaped micelle (left) and cartoon representation (middle) of compound 2-bound complex (**a**), compound 2 and GLP-1-bound complex (**b**), and compound 2 and LY3502970-bound complex (**c**). The chemical structures of compound 2 and LY3502970 are shown in the right panel of **a** and **c**. Compound 2 and GLP-1-bound GLP-1R in red; compound 2-bound GLP-1R in hot pink; compound 2 and LY3502970-bound GLP-1R in blue; GαS Ras-like domain in yellow; Gβ subunit in cyan; Gγ subunit in navy blue; Nb35 in gray; GLP-1 in green; compound 2 in orange; LY3502970 in purple.

### Binding of compound 2

The compound 2–GLP-1–GLP-1R–G_s_ complex has a typical active assembly close to that of GLP-1–GLP-1R–G_s_ with Cα root mean square deviation (RMSD) of 0.71Å for the whole complex (Fig. 2a). Compared with the peptide-free *apo* and intermediate state structures, compound 2 rendered the TM bundle and the ECD of GLP-1R undergo extensive conformational transitions (Supplementary Figs. 4 and 5). Specifically, starting from a closed conformation in *apo* state, the ECD of GLP-1R bound by either compound 2 or truncated peptide agonist (peptide 5)^10^ rotated clockwise and approached to ECL1 and ECL2. The extracellular parts of TM1 and TM2 moved inward to the center, while TM7 moved outwards accompanied by conformational changes of ECL1. In the intracellular domain, the sharp kink around the conserved Pro^6.47b^-X-X-Gly^6.50b^ motif in TM6 pivoted the intracellular half of TM6 to move outwards by 18.4Å (measured by Cα carbon of K346^6.35b^), to the same extent as that achieved by orthosteric agonists (Supplementary Fig. 4). This was joined by a modest outward movement of TM5 (3.6Å measured by Cα carbon of K336^5.66b^), thereby creating an intracellular crevice to accommodate G_s_ coupling^11,12,18^. The latter was anchored by the α5 helix of Gαs (GαH5) and formed rich contacts with TMs 2-7 and ICLs 1-3 (Supplementary Fig. 6).

**Fig. 2.**
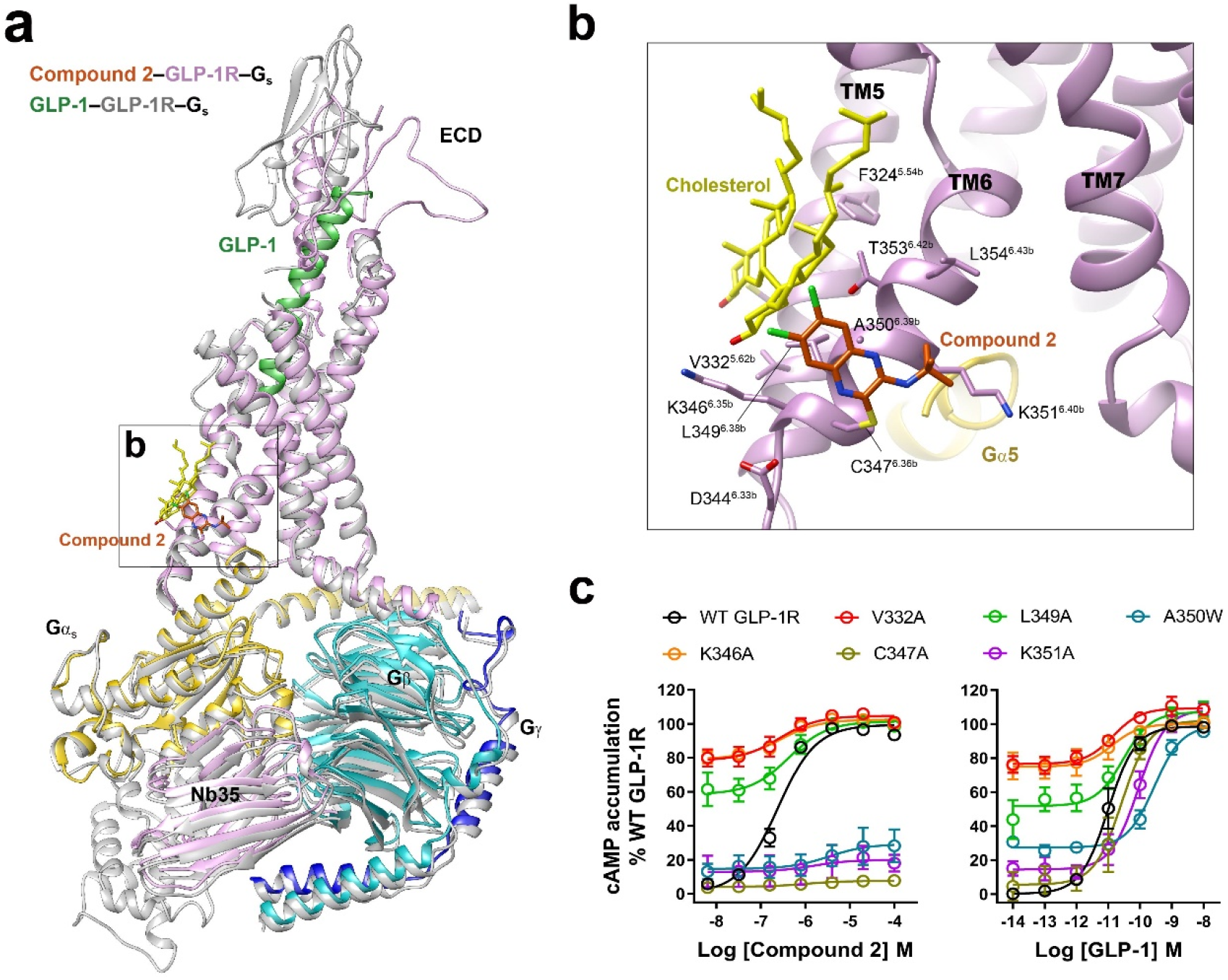
Unique agonist-binding site in GLP-1R for compound 2. **a**, Superimposition of the active state GLP-1R bound by compound 2 or GLP-1 (PDB code: 6X18) reveals a high structural similarity except the ECD region. **b**, Compound 2 covalently binds to C347^6.36b^ and mounts on the membranefacing surface of the cytoplasmic end of TM6. **c,** Representative mutation effects on the ago-allosterism associated with compound 2. Data are presented as means±S.E.M of three independent experiments. WT, wild-type. Source data are provided as a Source Data file.

Instead of occupying the orthosteric binding pocket where peptide or small molecule agonists generally bind, the high-resolution cryo-EM map demonstrates that compound 2 is covalently bonded to C347^6.36b^ (Wootten numbering in superscript^37^) and mounted on the membrane-facing surface of the cytoplasmic end of TM6, providing solid structural evidence of a unique binding site for the ago-allosteric modulator (Fig. 2b and Supplementary Fig. 3d). Such a covalent modification is supported by a previous report showing that non-sulfonic substituents at C-2 position (methylsulfone) failed to produce measurable cAMP responses^38^, consistent with our mutagenesis results, where C347^6.36b^A mutation diminished the potency of compound 2 without affecting that of GLP-1 (Fig. 2c and Supplementary Table 2). Compound 2 forms predominantly hydrophobic interactions with the adjacent residues in TM6. The tert-butyl moiety of compound 2 points to TM7 and makes hydrophobic contacts with A350^6.39b^ and K351^6.40b^, replacement at C-3 position by polar functional groups caused dramatic decline in its potency and efficacy^38^. The dichloroquinoxaline group extends to ICL3 forming nonpolar interactions with K346^6.35b^, C347^6.36b^ and a cholesterol molecule in TM6, introduction of electron-donating substituents or replacement of quinoxalines by benzimidazole or quinoline at C-6 and C-7 positions led to poor tolerance^38^. The bulky A350^6.39b^W and K351^6.40b^A mutants almost abolished the maximal response (Emax) of GLP-1R-mediated cAMP accumulation in presence of compound 2, while V332^5.62b^A, K346^6.35b^A and L349^6.38b^A mutants greatly elevated basal cAMP activities but significantly diminished the efficacy of the response upon stimulation by either compound 2 or GLP-1, suggesting these mutations affect the kink of TM6 required for receptor activation. The same covalent mechanism of action was previously demonstrated for another PAM of GLP-1R (i.e., BETP), showing that in addition to C347, BETP also formed a covalent adduct with C438 in the C terminus^27^ that is invisible in the current GLP-1R complex structures. Although mutation of C438 did not alter the PAM activity of BETP or compound 2 (Ref^27^), it cannot rule out possible formation of a covalent adduct at C438 in our compound 2-bound GLP-1R structure.

### ECD conformation

The most profound structural feature in the extracellular half of compound 2-bound GLP-1R is the unique position and orientation of ECD, distinct from all available full-length GLP-1R structures reported to date (Fig. 3). The tripartite α-β-β/α architecture of ECD observed in *apo* or peptide-bound structures^39^ was partially disturbed in the compound 2–GLP-1R–G_s_ complex except for the N-terminal α-helix (residues 29 to 49). GLP-1-bound ECD was shown to be fully extended^11,34^ whereas that of compound 2-bound folded down towards the TMD core and penetrated into the orthosteric binding pocket through its N-terminal α-helix and loop by inward movement of 9.0Å (measured at the Cα of R40). Notably, the orientation of ECD is distinct from that of GLP-1-bound GLP-1R with a rotation angle of 97.4 degrees: the former pointed from ECL2 to TM1-TM2 and the latter oriented from ECL1 to TM5-TM6 (Fig. 3a). The tip of the N-terminal α-helix (T29) moved by 19.9Å and inserted into a cleft between TM1 and TM2, partially overlapping with the recently reported allosteric site of PAM LSN3160440 (ref. ^17^). Locked by the conserved disulfide bond (C46-C71) with the N-terminal α-helix, β1 strand (residues 61 to 77) of GLP-1R ECD also shifted towards ECL1 by 9.3Å (measured at the Cα of E68). Such a movement consequently shortened the length of α-helix at ECL1 in *apo* state and extended TM2 by one turn, thereby stabilizing the ECD conformation via an extensive network of complimentary polar and non-polar contacts (Fig. 3a and Supplementary Fig. 5). Consistently, our molecular dynamic (MD) simulations found that the ECD interacted with the TMD intimately, such that the N-terminal α-helix stably inserted to the TMD core (Supplementary Fig. 7).

**Fig. 3.**
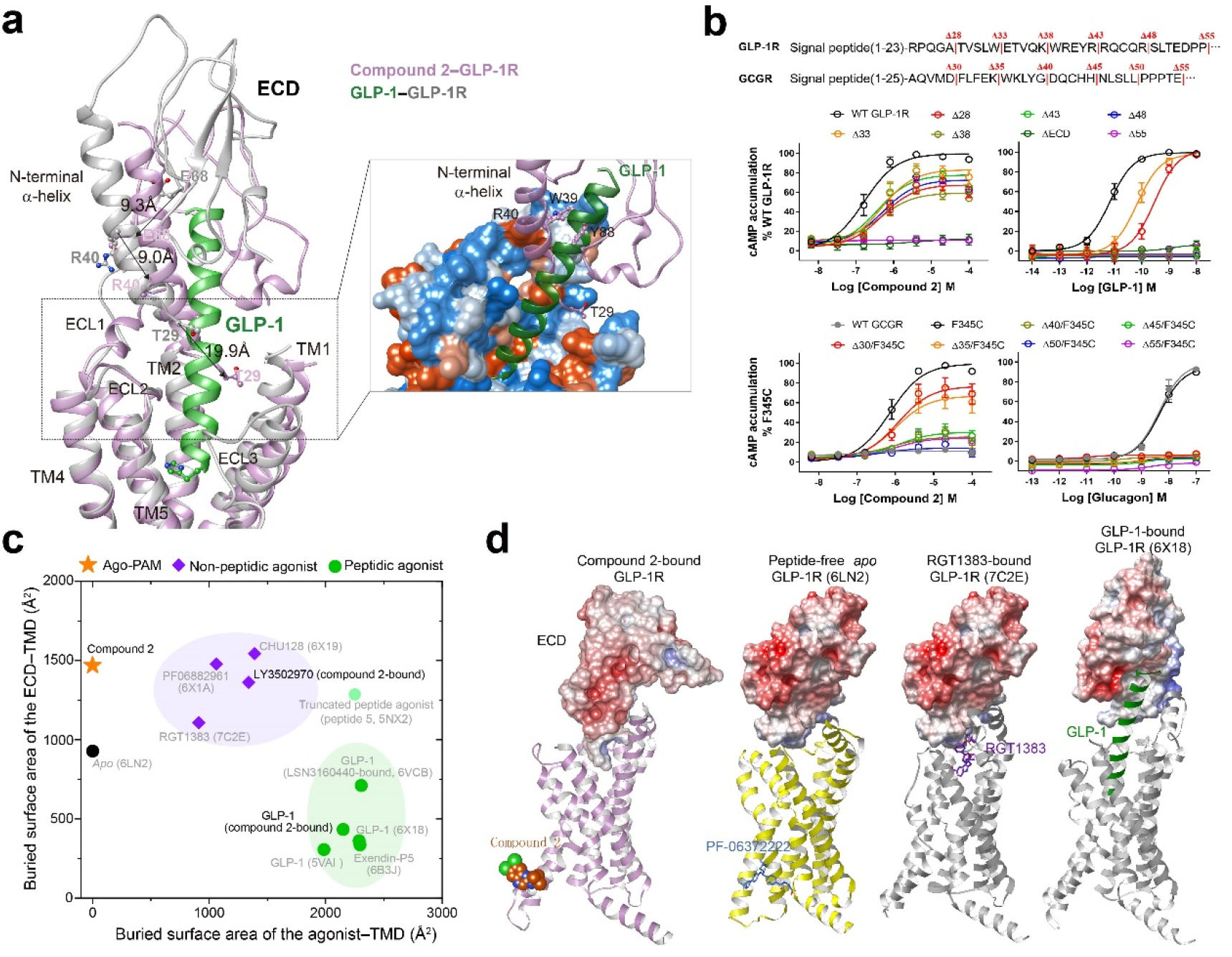
Diversified ECD engagements in GLP-1R activation. **a,** Comparison of the ECD conformation between the GLP-1-bound (PDB code: 6X18) and compound 2-bound GLP-1R. Close-up view of the interaction between ECD and TMD shows that the N-terminal α-helix of ECD penetrated into the orthosteric TMD pocket by a distinct orientation. **b,** Signaling profiles of ECD-truncated GLP-1R and GCGR in response to their cognate peptides (GLP-1 and glucagon) and compound 2. Compound 2 is verified as an agonist for GCGR mutant F345^6.36b^C. Data are presented as means±S.E.M of three independent experiments. WT, wild-type. Δ, residue truncation. **c,** Scatter plot of TMD-ECD/agonist interaction of all available GLP-1R structures with visible ECDs. X and Y axis represent the buried interface area of agonist-TMD and ECD-TMD, respectively. The interface areas were calculated using freeSASA. **d,** Surface representation of diversified conformations of ECD in representative GLP-1R structures. Source data are provided as a Source Data file.

To reveal functional roles of the ECD in the presence of either compound 2 or GLP-1, we truncated the N-terminal α-helix in a systemic manner and measured cAMP responses subsequently (Fig. 3b and Supplementary Table 2). For compound 2, truncation of a large portion (28-48 residues) of the N-terminal α-helix resulted in a dramatic efficacy decrease of compound 2, implying the role of the tip of N-terminal α-helix in the ago-allosterism. Removal of the N-terminal 55 residues or ECD deletion completely abolished the activation effect of compound 2 despite that it binds to the cytoplasmic side of TM6, consistent with our previous report that ECD is required for GLP-1R and GCGR activation^34^. Besides, similar dose-response characteristics between GLP-1 and compound 2 suggest that the latter is capable of stimulating GLP-1R independently (ago-allosteric activation). In contrast, the action of GLP-1 is dependent on the ECD which binds to the C-terminal half of the peptide. Truncation of the ECD by 28 or 33 residues reduced GLP-1 potency by 50- and 13-fold, respectively. In line with the previous study^34^, cAMP signaling was completely abolished when 38 residues were truncated, but this shortened construct still worked for compound 2. This phenomenon was also observed in constructs expressing wild-type and mutant glucagon receptors. For instance, F345^6.36b^C mutant of GCGR was sufficient to confer sensitivity to compound 2 (Fig. 3b and Supplementary Table 3). These results show that the ECD requirement is ligand-dependent and is differentiated between orthosteric and allosteric modulators.

To further address this point, we analyzed the TMD-ECD interacting patterns by measuring the buried surface area between TMD and ECD or between TMD and agonists for all available GLP-1R structures with visible ECDs (Fig. 3c, d). In the *apo* state, its ECD adopted a closed TMD-interacting conformation with a buried surface area of 928Å^2^. When bound by peptide agonists such as GLP-1 and exendin-P5, their ECDs stood up along the α-helical peptide and made extensive contacts with the C-terminal half of the peptide, thereby limiting the contact with TMD (~350Å^2^). Meanwhile, the N-terminal half of peptide inserted into the TMD core with massive contacts (buried surface area of more than 2000Å^2^). As a comparison, small molecules showed much reduced TMD-interacting surface area (~1000Å^2^). Complementally, its ECD folded down towards the TMD to stabilize the complex with a buried surface area of ~1500Å^2^, remarkably higher than that of peptidic ligands. In the case of compound 2-bound structure, the N-terminal α-helix of ECD penetrated to the TMD core and formed extensive contacts with residues in TM1-3, ECL1 and ECL2, resulting in an ECD-TMD interacting surface area of 1438.4Å^2^, similar to other small molecule agonists. The dynamic nature of ECD observed in this study is consistent with previous findings that GLP-1R is equipped with high versatility to recognize a wide range of ligands with distinct chemotypes^40,41^ and to participate in diversified receptor activation processes^13,34^ (Supplementary Fig. 4).

### Receptor activation

In spite of various conformational changes upon ligand binding, signaling initiation by either peptidic or small molecule agonists shares a common pathway, *i.e*., reorganization of the central polar network, HETX motif and TM2-6-7-helix 8 polar network, as well as the hallmark outward movement of TM6 (ref. ^11,13,42^). Thus, the complexes of GLP-1-GLP-1R-G_s_ and compound 2-GLP-1R-G_s_ displayed almost identical conformation reflecting the above polar network rearrangement and TM6 movement (Fig. 4). However, there are some distinct structural features in compound 2 alone bound receptor. At the bottom of the orthosteric pocket, the side chain of R310^6.40b^ pointed to the TMD core and formed a cation-pi stacking with Y241^3.44b^, where it pointed to ECL3 and made no contact with TMD core in the GLP-1 bound structure. Another notable difference resides in H363^6.52b^, which is adjacent to the conserved Pro^6.47b^-X-X-Gly^6.50b^ motif that pivots the intracellular half of TM6 to move outwards. Among all the active GLP-1R structures, H363^6.52b^ dipped into the TMD core whereas it is reoriented ~90° to an outside-facing position and formed a hydrogen bond with Q394^7.49b^ through dismissal of the hydrogen bond with the backbone carbonyl oxygen atom of P358^6.47b^. The distinct features manifested by R310^6.40b^ and H363^6.52b^ are likely to be responsible for the kink formation induced by compound 2, in line with previous mutagenesis results that mutation at these two sites altered GLP-1R signaling profiles (cAMP, pERK and iCa^2+^)^4,37^.

**Fig. 4.**
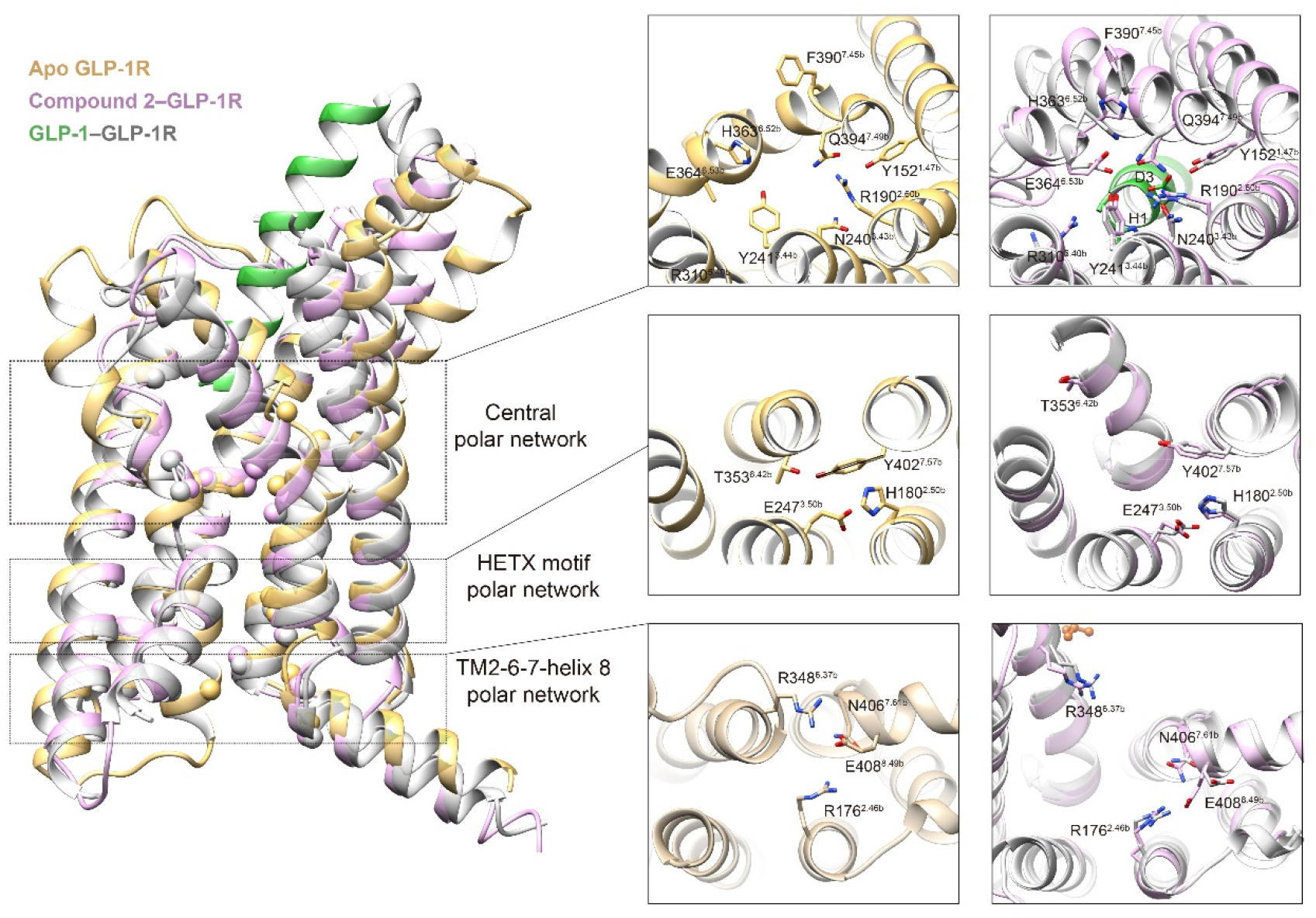
Comparison of the polar network rearrangement upon GLP-1R activation triggered by compound 2 and GLP-1. Left, superimposition of compound 2 bound GLP-1R with peptide-free *apo* state (PDB code: 6LN2) and GLP-1 bound active state (PDB code: 6X18). Upon receptor activation, three layers of polar network (central polar network located on the bottom of the orthosteric binding pocket, family-wide conserved HTEX motif polar network and TM2-6-7-helix 8 polar network that topologically close to G protein) were reorganized. The Cα atoms of the residues participated in the polar network rearrangements upon receptor activation are shown as spheres. Right, comparison of residual conformation between *apo* state, GLP-1-bound and compound 2-bound active states for three layers of polar network, whose residues are shown as stick with Wootten numbering in superscript.

As expected, the compound 2–GLP-1R–G_s_ complex exhibited a remarkable similarity to GLP-1-bound structures in terms of G protein-binding interface, consistent with a common mechanism of G_s_ protein engagement. Nonetheless, one additional contact was found to be unique for compound 2-bound structure, *i.e*., a hydrogen bond between the backbone atom of L260 in ICL2 and R38 in the GαH5 subunit (Supplementary Fig. 6).

### Positive allosterism

In line with previous findings^25^, we found that the potency of GLP-1was not changed but its binding affinity was enhanced by compound 2. The same phenomenon was observed for LY3502970 (Supplementary Fig. 8a). The high-resolution cryo-EM maps of the compound 2–GLP-1–GLP-1R–G_s_ and compound 2–LY3502970-GLP-1R–G_s_ complexes provide a good opportunity to further investigate compound 2-associated positive allosterism^25^. As shown in Fig. 5a, the overall structure of the compound 2–GLP-1–GLP-1R–G_s_ resembles that without this ago-PAM, with a Cα RMSD of 1.0Å. This observation also applies to LY3502970–GLP-1R–G_s_ with or without compound 2 (Cα RMSD of 0.8Å). The binding poses of compound 2 in these two structures are nearly identical to that of compound 2 alone except for a slightly different orientation (Fig. 5b and Supplementary Fig. 3d). As seen in the compound 2–GLP-1R–G_s_ complex, two cholesterol molecules were also found in the identical position of the compound 2–GLP-1–GLP-1R–G_s_ and compound 2–LY3502970–GLP-1R–G_s_ complexes, implying a common role for the two cholesterol molecules. In the presence of compound 2, both GLP-1 and the ECD moved inwards to the TMD core by 0.7 and 1.3Å (measured at the Cα of Y19 in GLP-1 and P90 in ECD), respectively. Upon binding of compound 2, several distinct contacts were formed including a hydrogen bond between T11 in GLP-1 and D372 in ECL3 as well as a salt bridge between R36 in GLP-1 and D215 in ECL1 (Supplementary Fig. 8b). Meanwhile, G_s_ moved upward by 0.5~0.8Å, especially the αN-helix in Gαs and Gβ subunits (Fig. 5c). Such an alteration may strengthen G protein coupling by introducing several newly formed polar interactions such as two salt bridges (R176^2.46b^ and E408^8.49b^, E423^8.64b^ and R46 in Gβ). Similar phenomenon was also observed in the compound 2–LY3502970–GLP-1R–G_s_ structure (Fig. 5c) and received the support of our MD studies in which compound 2-bound GLP-1R could stably bind to GLP-1 in the absence of G protein (Supplementary Fig. 9).

**Fig. 5.**
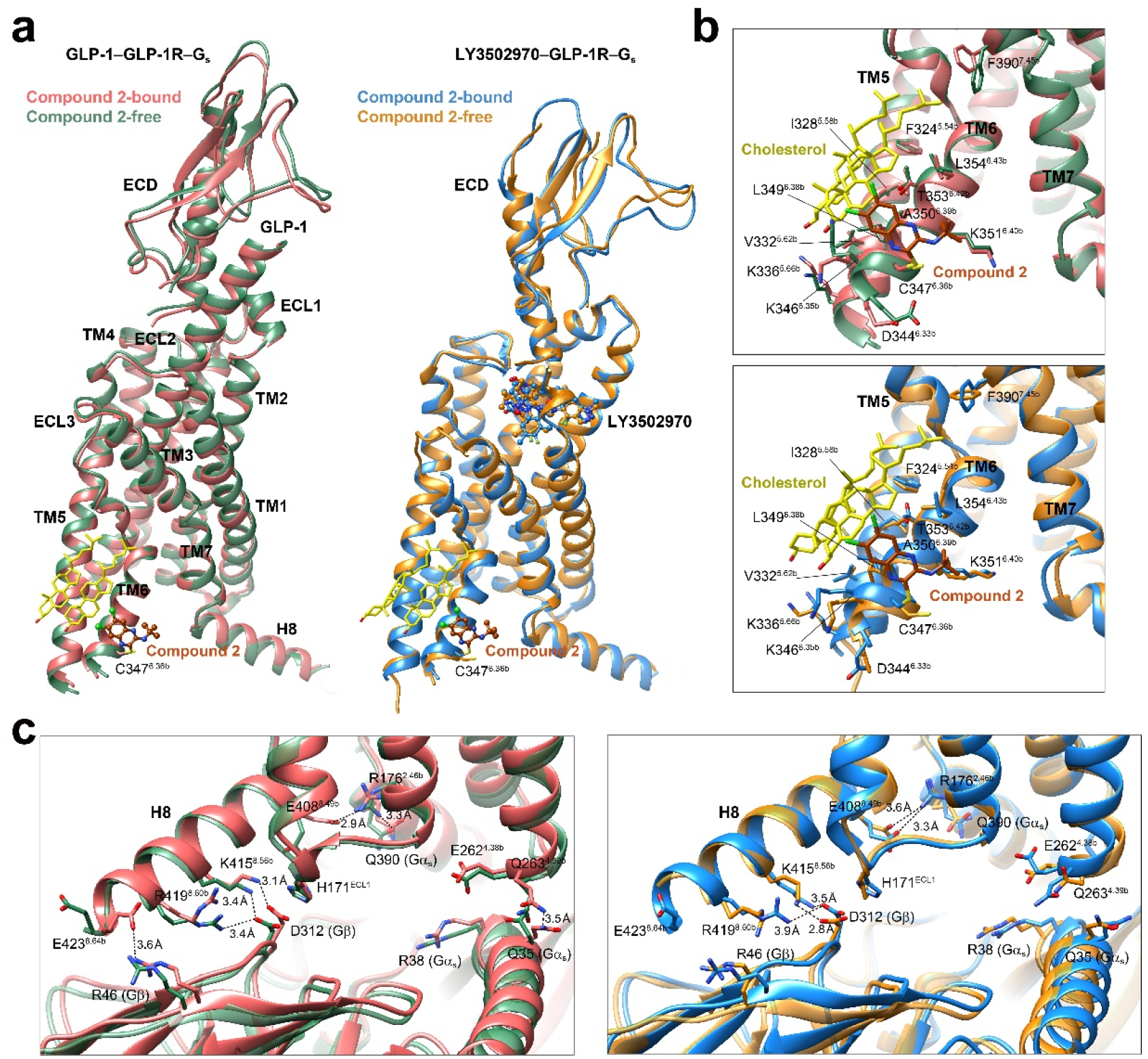
Structural insights into the ago-allosterism associated with compound 2. **a**, Superimposition of the GLP-1–GLP-1R–G_s_ (left, PDB code: 6X18) and LY3502970–GLP-1R–G_s_ (right, PDB code: 6XOX) structures with that bound to compound 2. G protein is omitted for clarity. **b**, Binding poses of compound 2. Two cholesterols bound to the TM5-TM6 cleft contribute hydrophobic interaction with compound 2. **c**, Comparison of the G protein-coupling interfaces of GLP-1 bound (left) and LY3502970 bound (right) active structures with that bound to compound 2.

## Discussion

It has been widely accepted that class B1 GPCRs adopt a two-step model for ligand binding and receptor activation^34,43^. However, this may not relevant to small molecule modulators that have different requirements for the ECD. Several recently determined non-peptidic ligand bound GLP-1R structures^13,14,16^ have uncovered previously unknown binding pockets that may have implications in their pharmacological profiles such as biased signaling and allosteric agonism (Supplementary Fig. 10). Ago-PAMs, acting as agonists on their own and as enhancers for orthosteric ligands, are capable of increasing agonist potency and providing additional efficacy^44^, thereby offering an alternative to conventional therapeutics. Mounted on the membrane-facing surface of the cytoplasmic tip of TM6, compound 2 was able to induce a newly discovered ECD conformation allowing its N-terminal helix to penetrate into the TMD core, a key step for the ago-allosterism observed experimentally, consistent with the intrinsic agonism hypothesis for the ECD of GLP-1R^40^. Interestingly, despite of the extracellular conformational differences induced by compound 2 and orthosteric agonists, their intracellular architectures underwent a similar reorganization of polar residues that enabled the formation TM6 sharp kink with the help of G protein binding^42,45^, indicative of GLP-1R activation (Fig. 6).

**Fig. 6.**
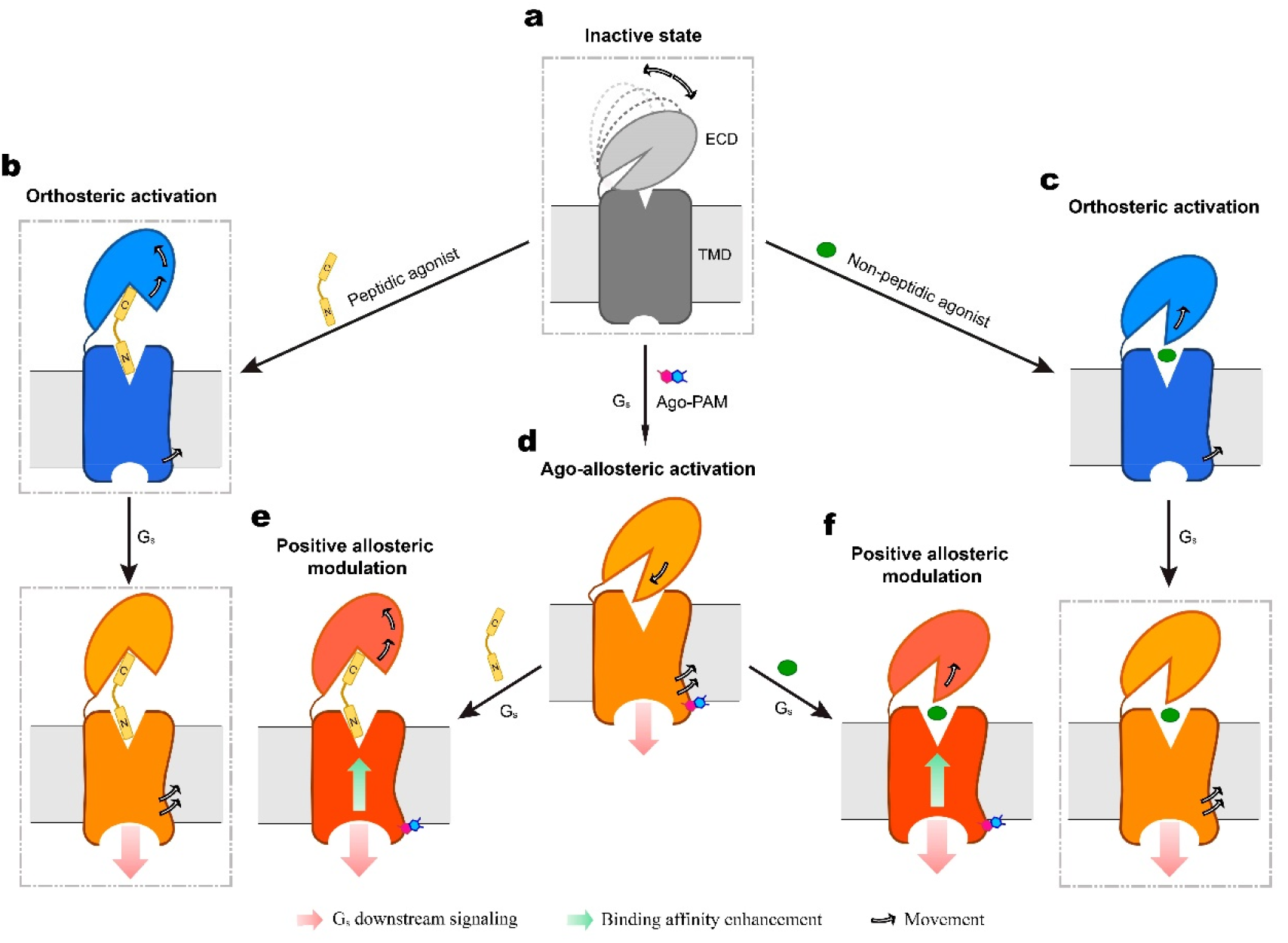
Postulated model of GLP-1R activation. **a**, In the absence of a ligand, the ECD of GLP-1R is dynamic with multiple conformations (dashed lines) but favors a closed state. Binding of peptidic (b) or non-peptidic (c) agonist triggers the ECD to disengage from the TMD thereby lowering the energy barrier of G protein coupling (**orthosteric activation**). **d**, Ago-PAM, such as compound 2, covalently binds to the cytoplasmic side of TM6 and induces its outward movement. Meanwhile, the ECD bends downwards to the TMD core thereby penetrating into the orthosteric pocket via the N-terminal α-helix. The TMD rearrangement, together with G_s_ binding, elict downstream signaling. (**ago-allosteric activation**). Ago-PAM also enhances both peptidic (e) and non-peptidic (f) agonist binding affinity (**positive allosteric modulation**). Inactive, intermediate (agonist-bound) and active (both agonist and G protein bound) conformations are colored in grey, blue and orange/orange-red, respectively. Movements of the ECD and the intracellular half of TM6 are indicated by arrows. Cell membranes are shown as grey bilayers. Previously reported GLP-1R structures are indicated by dashed-line boxes.

Diversified ECD conformations represent the dynamic nature of the receptor in response to a variety of external stimuli. In the peptide-free *apo* state, the ECD is in favor of a TMD closed conformation stabilized by ECL1 and ECL3 (ref. ^18^). Ligand binding triggers dissociation of the ECD from the TMD, allowing the peptide N-terminus to insert into the orthosteric binding pocket for receptor activation. Unlike peptide, the binding of compound 2 and other small molecule modulators require a complementary conformation of the ECD for ligand recognition, such that the N-terminal α-helix of the compound 2–GLP-1R–G_s_ deeply inserted to the orthosteric binding pocket as opposed to other GLP-1R structures (Fig. 6), suggesting its unique activation mechanism. In line with this view, we found that the ECD-TMD interaction is crucial for compound 2 and GLP-1 to function. However, this interaction was not required for small molecule agonist TT-OAD2 that elicited cAMP signaling without the presence of an ECD^14^. Reduction in cAMP responses to compound 2 following truncation of the N-terminal helix indicates that the ability of compound 2 to engender a stable ECD conformation may contribute to the “ago” effect on cAMP production. In line with the cryo-EM structure of compound 2-bound GLP-1R, the MD simulations indicate that the N-terminal helix consistently inserts to the TMD core thereby stabilizing the interaction between the ECD and ECL1. This implies that the insertion of the ECD N-terminal helix to the TMD core is a unique feature of compound 2-bound GLP-1R. Therefore, the ECD of GLP-1R plays a pivotal role in stabilizing receptor conformation and facilitating its activation as previously proposed^40,46^.

Collectively, our structures reveal an ago-PAM mechanism for GLP-1R, in which the rearrangement of conserved polar network is trigged by TM6 movement and stabilized by the ECD-TMD interaction. This work expands our understanding of GLP-1R activation and provides valuable insights into potential application of ago-allosterism in drug discovery.

## Methods

### Cell culture

*Spodoptera frugiperda* (*Sf9*) insect cells (Expression Systems) were grown in ESF 921 serum-free medium (Expression Systems) at 27°C and 120 rpm. HEK 293T cells (Cell Bank at the Chinese Academy of Sciences) were cultured in Dulbecco’s modified Eagle’s medium (DMEM, Life Technologies) supplemented with 10% fetal bovine serum (FBS, Gibco) and maintained in a humidified chamber with 5% CO_2_ at 37°C. CHO-K1 (ATCC #CCL-61) cells were maintained in F12 containing 10% (v/v) FBS, 100 units/mL penicillin and 100 mg/mL streptomycin at 37°C in 5% CO_2_.

### Constructs

The human GLP1-R was modified with its native signal sequence (M1-P23) replaced by the haemagglutinin (HA) signal peptide to facilitate receptor expression. To obtain a GLP1-R–G_s_ complex with good homogeneity and stability, we used the NanoBiT tethering strategy, in which the C-terminus of rat Gβ1 was linked to HiBiT subunit with a 15-amino acid polypeptide (GSSGGGGSGGGGSSG) linker and the C-terminus of GLP-1R was directly attached to LgBiT subunit followed by a TEV protease cleavage site and a double MBP tag (Supplementary Fig. 1a). A dominant-negative human Gαs (DNGαs) with 8 mutations (S54N, G226A, E268A, N271K, K274D, R280K, T284D and I285T) was generated by site-directed mutagenesis to limit G protein dissociation^47^. An engineered G_s_ construct (mini-G_s_) was used for expression and purification of the compound 2–LY3502970–GLP-1R complex, which was designed based on that used in the determination of A2AR-mini-G_s_ crystal structure^48^. The replacement of original Gαs α-helical domain (AHD, V65-L203) with that of human Gi1 (G60-K180) provided the binding site for Fab_G50, an antibody fragment used to stabilize the rhodopsin-G_i_ complex^49^. Additionally, substitution of N-terminal histidine tag (His6) and TEV protease cleavage site with the N-terminal eighteen amino acids (M1-M18) of human G_i1_ made this chimeric G_s_ capable of binding to scFv16 used to stabilize GPCR-G_i_ or -G_11_ complexes^50,51^. The constructs were cloned into both pcDNA3.1 and pFastBac vectors for functional assays in mammalian cells and protein expression in insect cells, respectively. All modifications of the receptor had no effect on ligand binding and receptor activation (Supplementary Fig. 1b and Supplementary Table 2). Other constructs including the fulllength and various N-terminal truncated human GLP-1R and GCGR were cloned into pcDNA3.1 vector for cAMP accumulation and whole cell binding assays (Supplementary Table 6).

### Formation and purification of complex

The Bac-to-Bac Baculovirus Expression System (Invitrogen) was used to generate high-titer recombinant baculovirus for GLP1R-LgBiT-2MBP, DNGαs, Gβ1-HiBiT and Gγ2. P0 viral stock was produced by transfecting 5 μg recombinant bacmids into *Sf9* cells (2.5 mL, density of 1 million cells per mL) for 96 h incubation and then used to produce P1 and P2 baculovirus. GLP1R-LgBiT-2MBP, DNGαs, Gβ1-HiBiT and Gγ2 were coexpressed at multiplicity of infection (MOI) ratio of 1:1:1:1 by infecting *Sf9* cells at a density of 3 million cells per mL with P2 baculovirus (viral titers>90%). Culture was harvested by centrifugation for 48 h post infection and cell pellets were stored at −80°C until use.

The cell pellets were thawed and lysed in a buffer containing 20 mM HEPES, pH 7.5, 100 mM NaCl, 10% (v/v) glycerol, 10 mM MgCl_2_, 1 mM MnCl_2_ and 100 μM TCEP supplemented with EDTA-free protease inhibitor cocktail (Bimake) by dounce homogenization. The complex formation was initiated by the addition of 10 μM GLP-1 or LY3502970 and/or 50 μM compound 2, 10 μg/mL Nb35 and 25 mU/mL apyrase (New England Bio-Labs). After 1.5 h incubation at room temperature (RT), the membrane was solubilized in the buffer above supplemented with 0.5% (w/v) lauryl maltose neopentyl glycol (LMNG, Anatrace) and 0.1% (w/v) cholesterol hemisuccinate (CHS, Anatrace) for 2 h at 4°C. The supernatant was isolated by centrifugation at 65,000 *g* for 30 min and incubated with amylose resin (New England Bio-Labs) for 2 h at 4°C. The resin was then collected by centrifugation at 500 *g* for 10 min and washed in gravity flow column (Sangon Biotech) with five column volumes of buffer containing 20 mM HEPES (pH 7.5), 100 mM NaCl, 10% (v/v) glycerol, 5 mM MgCl2, 1 mM MnCl2, 25 μM TCEP, 0.1% (w/v) LMNG, 0.02% (w/v) CHS, 2 μM GLP-1 or LY3502970 and/or 10 μM compound 2, followed by washing with fifteen column volumes of buffer containing 20 mM HEPES (pH 7.5), 100 mM NaCl, 10% (v/v) glycerol, 5 mM MgCl2, 1 mM MnCl2, 25 μM TCEP, 0.03% (w/v) LMNG, 0.01% (w/v) glyco-diosgenin (GDN, Anatrace), 0.008% (w/v) CHS, 2 μM GLP-1 or LY3502970 and/or 10 μM compound 2. The protein was then incubated overnight with TEV protease (customer-made) on the column to remove the C-terminal 2MBP-tag in the buffer above at 4°C. The flow through was collected next day and concentrated with a 100 kDa molecular weight cut-off concentrator (Millipore). The concentrated product was loaded onto a Superdex 200 increase 10/300 GL column (GE Healthcare) with running buffer containing 20 mM HEPES (pH 7.5), 100 mM NaCl, 10 mM MgCl2, 100 μM TCEP, 1 μM GLP-1 or LY3502970 and/or 5 μM compound 2, 0.00075% LMNG, 0.00025% GDN and 0.0002% (w/v) CHS. The fractions for monomeric complex were collected and concentrated to 15-20 mg/mL for electron microscopy examination.

### Expression and purification of Nb35

Nb35 with a C-terminal 6 × His-tag was expressed in the periplasm of *E. coli* BL21 (DE3), extracted and purified by nickel affinity chromatography^52^. Briefly, Nb35 target gene was transformed in BL21 and grown in TB culture medium with 100 μg/mL ampicillin, 2 mM MgCl2 and 0.1 % (w/v) glucose at 37°C, 180 rpm. The expression was induced by adding 1 mM IPTG when OD600 reached 0.7-1.2. The cell pellet was collected by centrifugation after overnight incubation at 28°C, 180 rpm and stored at −80°C until use. The HiLoad 16/600 Superdex 75 column (GE Healthcare) was used to separate the monomeric fractions of Nb35 with running buffer containing 20 mM HEPES, pH 7.5 and 100 mM NaCl. The purified Nb35 was flash frozen in 30% (v/v) glycerol by liquid nitrogen and stored in −80°C until use.

### Cryo-EM data acquisition

The concentrated sample (3.5 μL) was applied to glow-discharged holey carbon grids (Quantifoil R1.2/1.3, 300 mesh), and subsequently vitrified using a Vitrobot Mark IV (ThermoFisher Scientific) set at 100% humidity and 4°C. Cryo-EM images were collected on a Titan Krios microscope (FEI) equipped with Gatan energy filter and K3 direct electron detector and performed using serialEM. The microscope was operated at 300 kV accelerating voltage and a calibrated magnification of ×81000 corresponding to a pixel size of 1.045Å. The total exposure time was set to 7.2 s with intermediate frames recorded every 0.2 s, resulting in an accumulated dose of 80 electrons per Å^2^ with a defocus range of −1.2 to −2.2 μm. Totally, 3,609 images for compound 2–GLP-1R–G_s_, 5,536 images for compound 2–LY3502970–GLP-1R–G_s_ and 4,247 images for compound 2–GLP-1–GLP-1R–G_s_ complexes were collected.

### Image processing

Dose-fractionated image stacks were subjected to beam-induced motion correction using MotionCor2 v1.4.2 (ref. ^53^). A sum of all frames, filtered according to the exposure dose, in each image stack was used for further processing. Contrast transfer function (CTF) parameters for each micrograph were determined by Gctf v1.06 (ref. ^54^). Particle selection, 2D and 3D classifications were performed on a binned dataset with a pixel size of 2.09Å using RELION-3.0.8-beta2 (ref. ^55^).

For the compound 2–GLP-1–GLP-1R–G_s_ complex, auto-picking yielded 1,186,340 particle projections were subjected to 3D classification with mask on the receptor to discard false positive particles or particles categorized in poorly defined classes, producing 703,620 particle projections for further processing. This subset of particle projections was subjected to further 3D autorefinement with mask on the complex, which were subsequently subjected to a round of 3D classifications with mask on the ECD. A selected subset containing 614,978 projections was subsequently subjected to 3D refinement with mask on the complex and Bayesian polishing with a pixel size of 1.045Å. After last round of refinement, the final map has an indicated global resolution of 2.5Å at a Fourier shell correlation (FSC) of 0.143. Local resolution was determined using the Bsoft package with half maps as input maps^56^.

For the compound 2–GLP-1R–G_s_ complex, auto-picking yielded 2,366,210 particles. Among them, 21.23% presented better densities on the ECD than other classifications. Thus, this subset was subjected to 3D classification with mask focused on the ECD and the TMD. Then 285,885 particles were selected for further 3D classification with mask focused on the ECD. Finally, 147,173 particles were subjected to 3D refinement with mask focused on the complex and Bayesian polishing with a pixel size of 1.045Å. The final map has an indicated global resolution of 3.3Å at a FSC of 0.143. Apart from that, there were 63.42% of the 2,366,210 particles holding better TMD and G protein densities. They were used for 3D classification with mask focused on the TMD and the G protein. Of these, 848,918 particles were selected for 3D classification focused on the TMD. Finally, 340,501 particles were used for final 3D refinement with mask focused on the complex and Bayesian polishing with a pixel size of 1.045Å. The final map has an indicated global resolution of 2.5Å at a FSC of 0.143.

For the compound 2-LY3502970–GLP-1R–G_s_ complex, CTF parameters were estimated with CTFFIND43. A total of 6,566,177 particles were automatically picked from 5552 images. Particle selection, 2D classification, 3D classification and refinement were performed using RELION-3.0.8-beta2. A data set of 345,411 particles was subjected to 3D refinements, yielding a final map with a global nominal resolution at 2.9Å by the 0.143 criteria of the gold-standard FSC. Half-reconstructions were used to determine the local resolution of each map.

### Model building and refinement

The structures of the LSN3160440–GLP-1–GLP-1R–G_s_ (PDB: 6VCB), GLP-1–GLP1R–G_s_ (PDB: 6X18) and RGT1383–GLP-1R–G_s_ (PDB: 6B3J) were used as an initial template for model building of compound 2, compound 2 plus GLP-1, and compound 2 plus LY3502970 bound complexes, respectively. Lipid coordinates and geometry restraints were generated using Phenix 1.16. Models were docked into the EM density map using UCSF Chimera 1.13.1. This starting model was then subjected to iterative rounds of manual adjustment and automated refinement in Coot 0.9.4.1 and Phenix 1.16, respectively. The final refinement statistics were validated using the module comprehensive validation (cryo-EM) in Phenix 1.16. Structural figures were prepared in UCSF Chimera 1.13.1 and PyMOL 2.1 (https://pymol.org/2/). The final refinement statistics are provided in Supplementary Table 1.

### cAMP accumulation assay

Peptide or small molecules stimulated cAMP accumulation was measured by a LANCE Ultra cAMP kit (PerkinElmer). Briefly, after 24 h transfection with various constructs, HEK 293T cells were digested by 0.2% (w/v) EDTA and washed once with Dulbecco’s phosphate buffered saline (DPBS). Cells were then resuspended with stimulation buffer (Hanks’ balanced salt solution (HBSS) supplemented with 5 mM HEPES, 0.5 mM IBMX and 0.1% (w/v) BSA, pH 7.4) to a density of 0.6 million cells per mL and added to 384-well white plates (3000 cells per well). Different concentrations of ligand in stimulation buffer were added and the stimulation lasted for 40 min at RT. The reaction was stopped by adding 5 μL Eu-cAMP tracer and ULight-anti-cAMP. After 1 h incubation at RT, the plate was read by an Envision plate reader (PerkinElmer) to measure TR-FRET signals (excitation: 320 nm, emission: 615 nm and 665 nm). A cAMP standard curve was used to convert the fluorescence resonance energy transfer ratios (665/615 nm) to cAMP concentrations.

### Whole cell binding assay

CHO-K1 cells were seeded to 96-well plates (PerkinElmer) coated with Fibronectin (Corning) at a density of 30,000 cells per well and incubated overnight. After 24 h transfection, cells were washed twice and incubated with blocking buffer (F12 supplemented with 33 mM HEPES, and 0.1% (w/v) BSA, pH 7.4) for 2 h at 37°C. Then, radiolabeled ^125^I-GLP-1 (40 pM, PerkinElmer) and increasing concentrations of unlabeled peptide were added and competitively reacted with the cells in binding buffer (PBS supplemented with 10% (w/v) BSA, pH 7.4) at RT for 3 h. After that, cells were washed with ice-cold PBS and lysed by 50 μL lysis buffer (PBS supplemented with 20 mM Tris-HCl and 1% (v/v) Triton X-100, pH 7.4). Finally, 150 μL of scintillation cocktail (OptiPhase SuperMix, PerkinElmer) was employed and radioactivity (counts per minute, CPM) determined by a scintillation counter (MicroBeta2 plate counter, PerkinElmer).

### Molecular dynamics simulation

Molecular dynamic simulations were performed by Gromacs 2018.5. The receptor was prepared and capped by the Protein Preparation Wizard (Schrodinger 2017-4), while the titratable residues were left in their dominant state at pH 7.0. The complexes were embedded in a bilayer composed of 167 POPC lipids, 42 cholesterols and solvated with 0.15M NaCl in explicitly TIP3P waters using CHARMM-GUI Membrane Builder v3.2.2 (ref. ^57^). The CHARMM36-CAMP force filed^58^ was adopted for protein, peptides, lipids and salt ions. Compound 2 was modelled with the CHARMM CGenFF small-molecule force field, program version 1.0.0 (ref. ^59^). The Particle Mesh Ewald (PME) method was used to treat all electrostatic interactions beyond a cutoff of 10Å and the bonds involving hydrogen atoms were constrained using LINCS algorithm^60^. The complex system was firstly relaxed using the steepest descent energy minimization, followed with slow heating of the system to 310 K with restraints. The restraints that adopted from the default setting in the CHARM-GUI webserver v3.2.2 (ref. ^57^) were reduced gradually over 50 ns, with a simulation step of 1 fs (see Supplementary Table 5 for more details). Finally, restrain-free production run was carried out for each simulation, with a time step of 2 fs in the NPT ensemble at 310 K and 1 bar using the Nose-Hoover thermostat and the semi-isotropic Parrinello-Rahman barostat^61^, respectively. The interface areas of agonist-TMD and ECD-TMD were calculated using FreeSASA 2.0 (ref. ^62^) using the Sharke-Rupley algorithm with a probe radius of 1.2Å.

### Statistical analysis

All functional study data were analyzed using Prism 8.0 (GraphPad) and presented as means ± S.E.M. from at least three independent experiments. Concentration-response curves were evaluated with a three-parameter logistic equation. The significance was determined with either two-tailed Student’s *t*-test or one-way ANOVA, and P<0.05 was considered statistically significant.

### Reporting summary

Further information on research design is available in the Nature Research Reporting Summary linked to this article.

## Data availability

All relevant data are available from the corresponding authors upon reasonable request. A reporting summary for this article is available as a Supplementary Information file. The raw data underlying Figs. 2c, 3b and Supplementary Figs. 1b-e, 5c, 8a are provided as a Source Data file. The atomic coordinates and electron microscopy maps have been deposited in the Protein Data Bank (PDB) under accession codes: 7DUR (compound 2–GLP–1R–G_s_ complex), 7DUQ (compound 2–GLP-1-GLP-1R–G_s_ complex) and 7E14 (compound 2–LY3502970–GLP-1R–G_s_ complex) and Electron Microscopy Data Bank (EMDB) accession codes: EMD-30867 (compound 2–GLP–1R-G_s_ complex), EMD-30886 (compound 2–GLP-1–GLP-1R–G_s_ complex) and EMD-30936 (compound 2-LY3502970–GLP-1R–G_s_ complex), respectively.

## Acknowledgements

We thank D.P. Yuan, W. Guo, F. Liu, B. Qiu, P.F. Lan and M. Lei for technical advice. This work was partially supported by National Natural Science Foundation of China 81872915 (M.-W.W.), 82073904 (M.-W.W.), 81922071 (Y.Z.), 81773792 (D.Y.), 81973373 (D.Y.) and 21704064 (Q.Z.); National Science & Technology Major Project of China–Key New Drug Creation and Manufacturing Program 2018ZX09735–001 (M.-W.W.) and 2018ZX09711002–002–005 (D.Y); the National Key Basic Research Program of China 2018YFA0507000 (M.-W.W.) and 2019YFA0508800 (Y.Z.), Shanghai Municipal Science and Technology Major Project 2019SHZDZX02 (H.E.X.); Ministry of Science and Technology of China Major Project XDB08020303 (H.E.X.); Shanghai Municipal Science and Technology Commission grant 19ZR1467500 (H.M.); Zhejiang Province Science Fund for Distinguished Young Scholars LR19H310001 (Y.Z.); Fundamental Research Funds for Central Universities 2019XZZX001-01-06 (YZ.); Novo Nordisk-CAS Research Fund grant NNCAS-2017–1-CC (D.Y.); The Young Innovator Association of CAS Enrollment (H.M. and L.H.Z.) and SA-SIBS Scholarship Program (L.H.Z. and D.Y). The cryo-EM data were collected at Cryo-Electron Microscopy Research Center, Shanghai Institute of Materia Medica.

## Author contributions

Z.T.C. and H.L.M. designed the expression constructs, purified the receptor complexes, screened specimen, prepared the final samples for negative staining/data collection towards the structure and participated in manuscript preparation; L.-N.C., H.L.M. and X.YZ. performed map calculation, structure analysis, figure preparation and participated in manuscript writing; Q.T.Z. performed structural analysis, MD simulations, figure preparation and participated in manuscript writing; Q.L. synthesized compound 2; Z.T.C. and A.T.D. conducted ligand binding and signaling experiments; C.Y.Y. and X.W. assisted in complex purification; T.X. assisted in structural analysis; L.H.Z. and P.Y.X. helped construct design and data analysis; W.H., X.Q.S. and W.G.Z. supplied LY3502970 and helped LY3502970 related structure determination; D.Y. supervised mutagenesis and signaling experiments, participated in data analysis and manuscript preparation; H.E.X., Y.Z., W.Z., and M.-W.W. initiated the project, supervised the studies, analyzed the data and wrote the manuscript with inputs from all co-authors.

## Additional information

**Supplementary information** is available for this paper at XXXX.

Correspondence and requests for materials should be addressed to D.Y., H.E.X., Y.Z. or M.-W.W.

## Supplementary Information

Brief description of what this file includes:

Supplementary Fig. 1 Functional validation of the receptor constructs and purification of the complexes.

Supplementary Fig. 2 Cryo-EM data processing and validation.

Supplementary Fig. 3 Near-atomic resolution model of the complexes in the cryo-EM density maps.

Supplementary Fig. 4 Comparison of available GLP-1R structures.

Supplementary Fig. 5 Unique ECD conformation in the compound 2-bound GLP-1R.

Supplementary Fig. 6 Comparison of G protein coupling between compound 2-bound and GLP-1-bound active GLP-1R in complex with G_s_.

Supplementary Fig. 7 Molecular dynamics (MD) simulations of compound 2-bound active GLP-1R.

Supplementary Fig. 8 Potentiation of GLP-1 and LY3502970 activity by compound 2.

Supplementary Fig. 9 MD simulations of compound 2–GLP-1–GLP-1R.

Supplementary Fig. 10 List of small molecule GLP-1R modulators with available structures.

Supplementary Table. 1 Cryo-EM data collection, refinement and validation statistics.

Supplementary Table. 2 *In vitro* pharmacology of GLP-1R.

Supplementary Table. 3 *In vitro* pharmacology of GCGR.

Supplementary Table. 4 Effects of residue mutation in the ECD-ECL1 interface on cAMP accumulation.

Supplementary Table. 5 Details of restraints applied during MD simulations.

Supplementary Table. 6 Primers used in this study.

**Supplementary Fig. 1.**
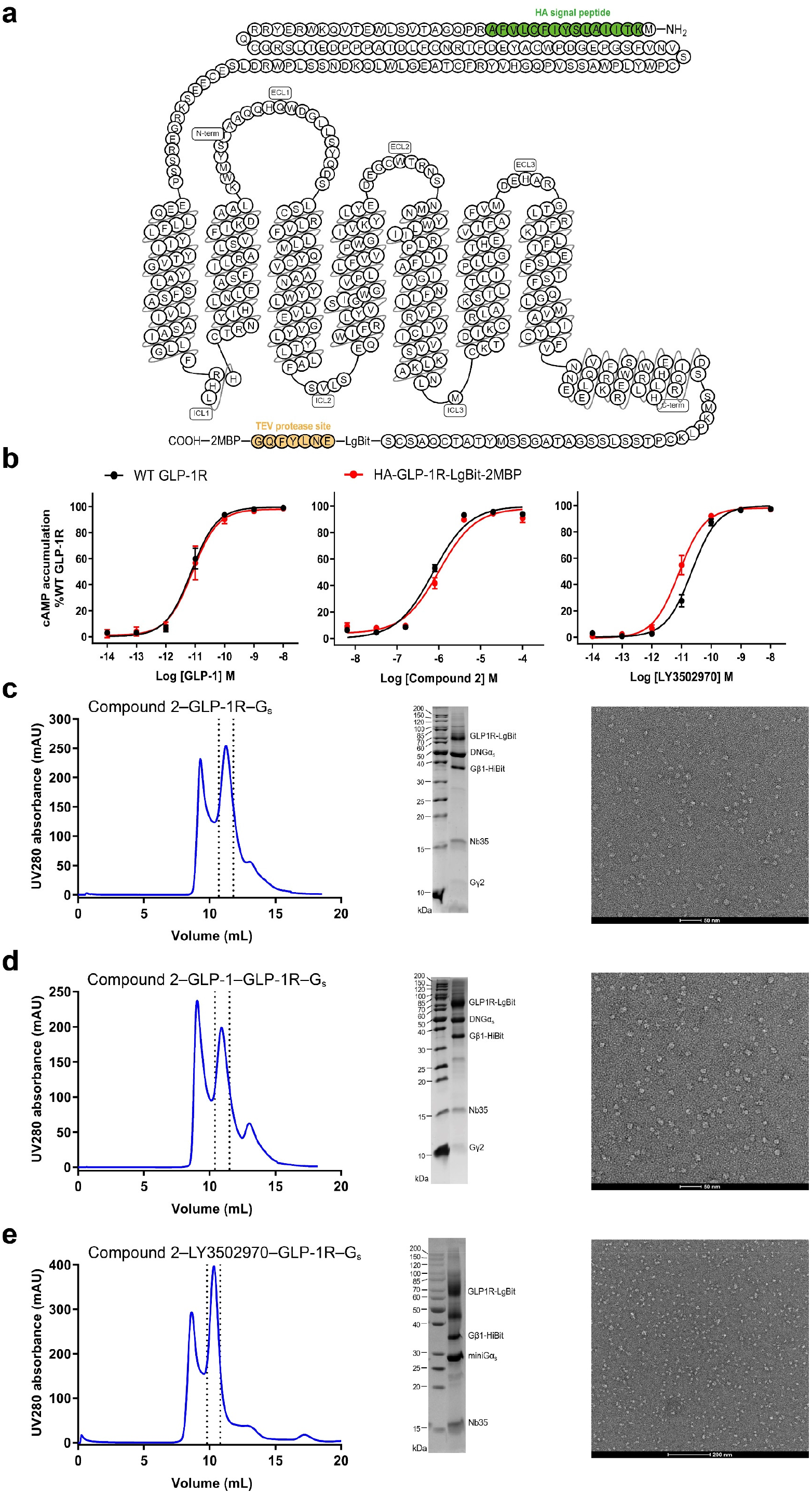
Functional validation of the receptor constructs and purification of the complexes. **a**, Schematic diagram of the receptor constructs used for structure determination. **b**, GLP-1, compound 2 and LY3502970 induced cAMP accumulation. Data are shown as means ± S.E.M from three independent experiments performed in technical triplicate. **c-e**, Analytical sizeexclusion chromatography, SDS-PAGE/Coomassie blue stain and representative negative staining image of the purified complexes: compound 2–GLP-1R–G_s_ complex (**c**), compound 2–GLP-1–GLP-1R–G_s_ complex (**d**) and compound 2–LY3502970–GLP-1R–G_s_ complex (**e**). These experiments were repeated independently twice with similar results. WT, wild-type. Source data are provided as a Source Data file.

**Supplementary Fig. 2.**
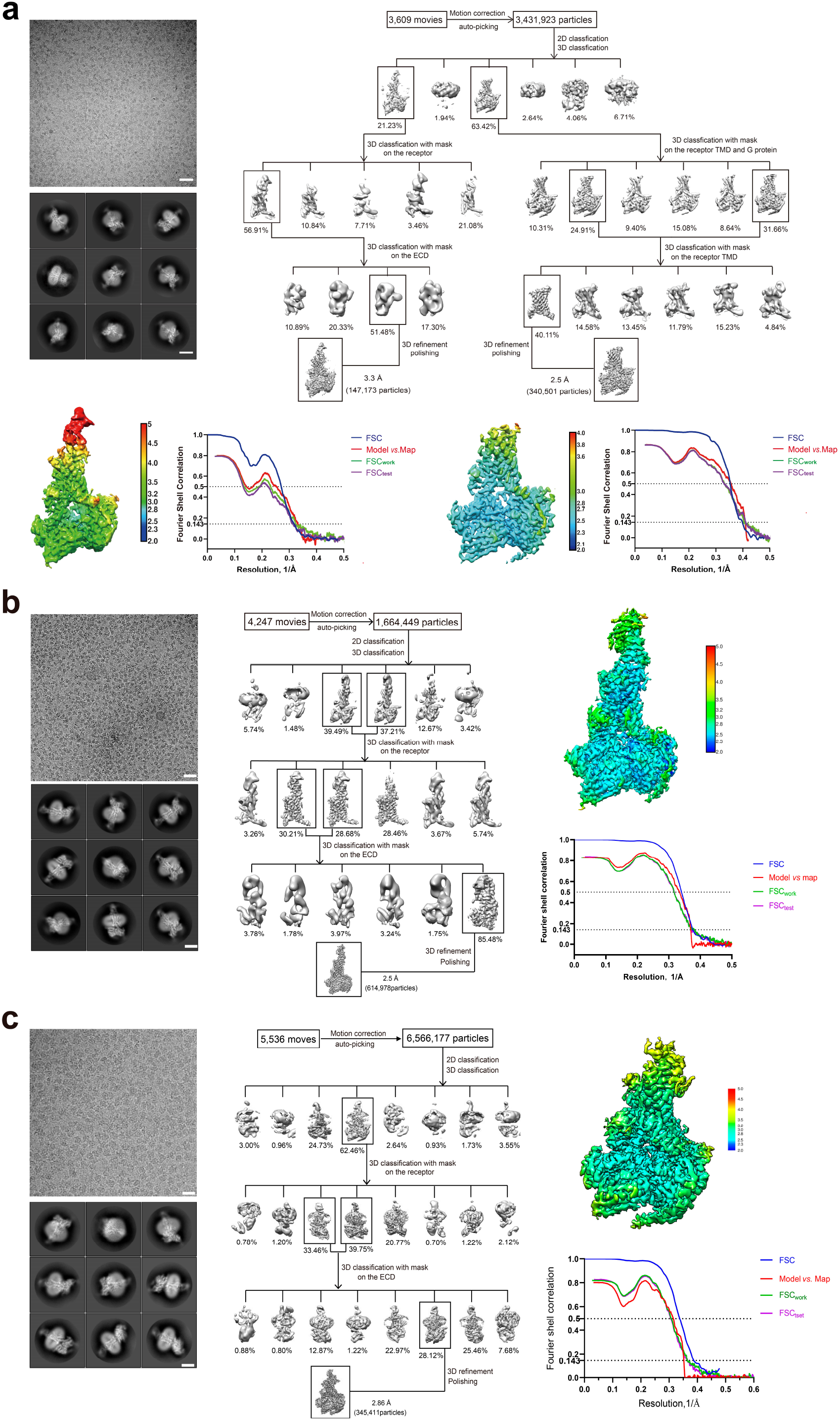
Cryo-EM data processing and validation. **a**, Compound 2–GLP-1R–G_s_ complex: top left, representative cryo-EM micrograph (scale bar: 40 nm) and two-dimensional class averages (scale bar: 5 nm); top right, flow chart of cryo-EM data processing; bottom left, local resolution distribution map of the complex with the ECD and Gold-standard Fourier shell correlation (FSC) curves of overall refined receptor; bottom right, local resolution distribution map of the complex without the ECD and FSC curves of overall refined receptor. **b**, Compound 2–GLP-1–GLP-1R–G_s_ complex: left, representative cryo-EM micrograph (scale bar: 40 nm) and twodimensional class averages (scale bar: 5 nm); middle, flow chart of cryo-EM data processing; right, local resolution distribution map of the complex and FSC curves of overall refined receptor. **c**, Compound 2–LY3502970–GLP-1R–G_s_ complex: left, representative cryo-EM micrograph (scale bar: 20 nm) and two-dimensional class averages (scale bar: 5 nm); middle, flow chart of cryo-EM data processing; right, local resolution distribution map of the complex and FSC curves of overall refined receptor. These experiments were repeated independently twice with similar results.

**Supplementary Fig. 3.**
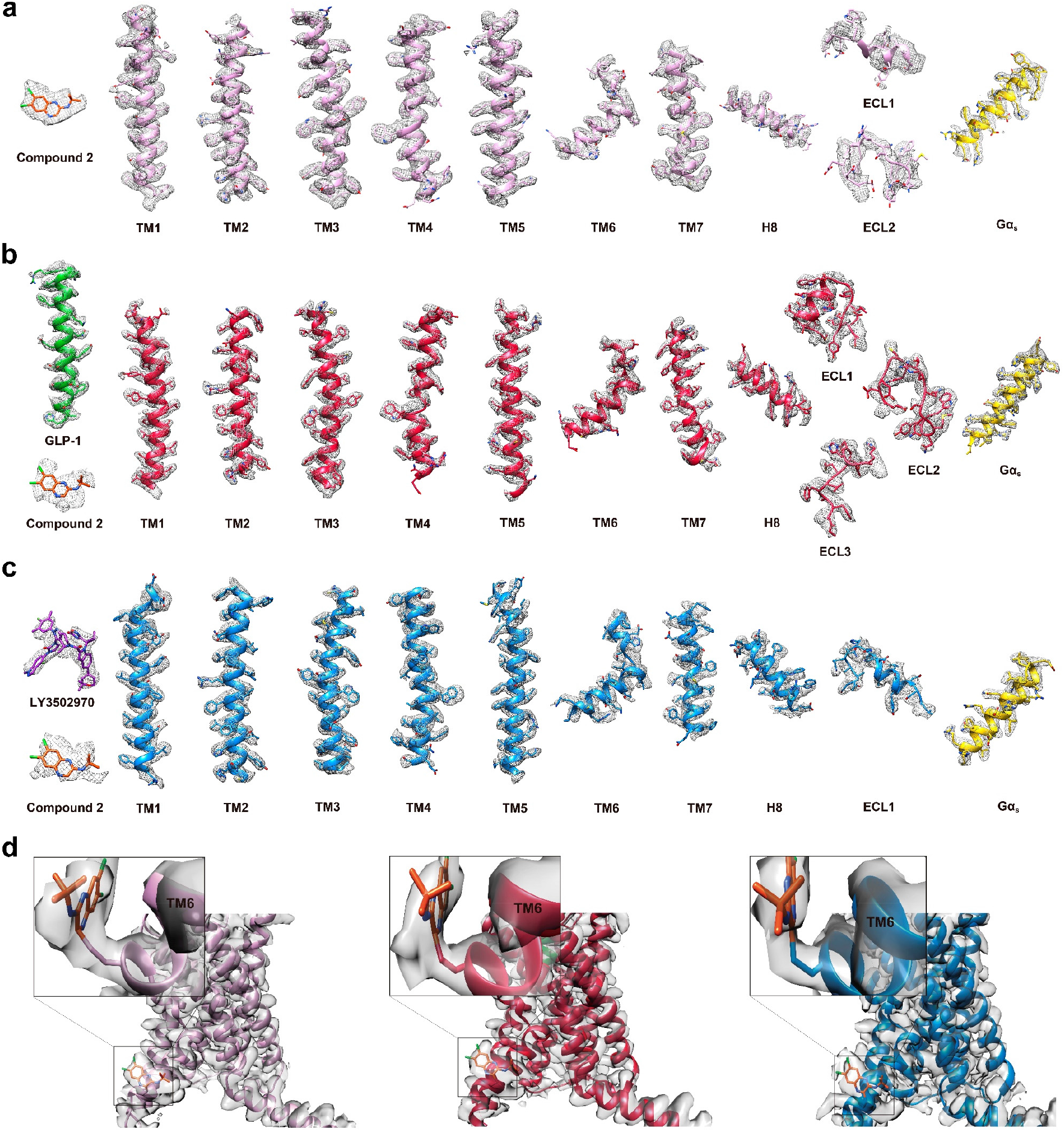
Near-atomic resolution model of the complexes in the cryo-EM density maps. **a**, EM density map and model of the compound 2–GLP-1R–G_s_ complex are shown for all seven-transmembrane (7TM) α-helices, helix 8 and all extracellular loops of GLP-1R, compound 2 and the α5-helix of the Gα_s_ Ras-like domain. **b**, EM density map and model of the compound 2–GLP-1–GLP-1R–G_s_ complex are shown for all 7TM α-helices, helix 8 and all extracellular loops of GLP-1R, GLP-1, compound 2 and the α5-helix of the Gαs Ras-like domain. **c**, EM density map and model of the compound 2–LY3502970–GLP-1R–G_s_ complex are shown for all 7TM α-helices, helix 8 and ECL1 of GLP-1R, compound 2, LY3502970 and the α5-helix of the Gαs Ras-like domain. **d**, The zoomed-in views of compound 2 and the interacting region of TM6 placed inside the density maps of compound 2–GLP-1 (left), compound 2–GLP-1–GLP-1R (middle) and compound 2–LY3502970–GLP-1R (right).

**Supplementary Fig. 4.**
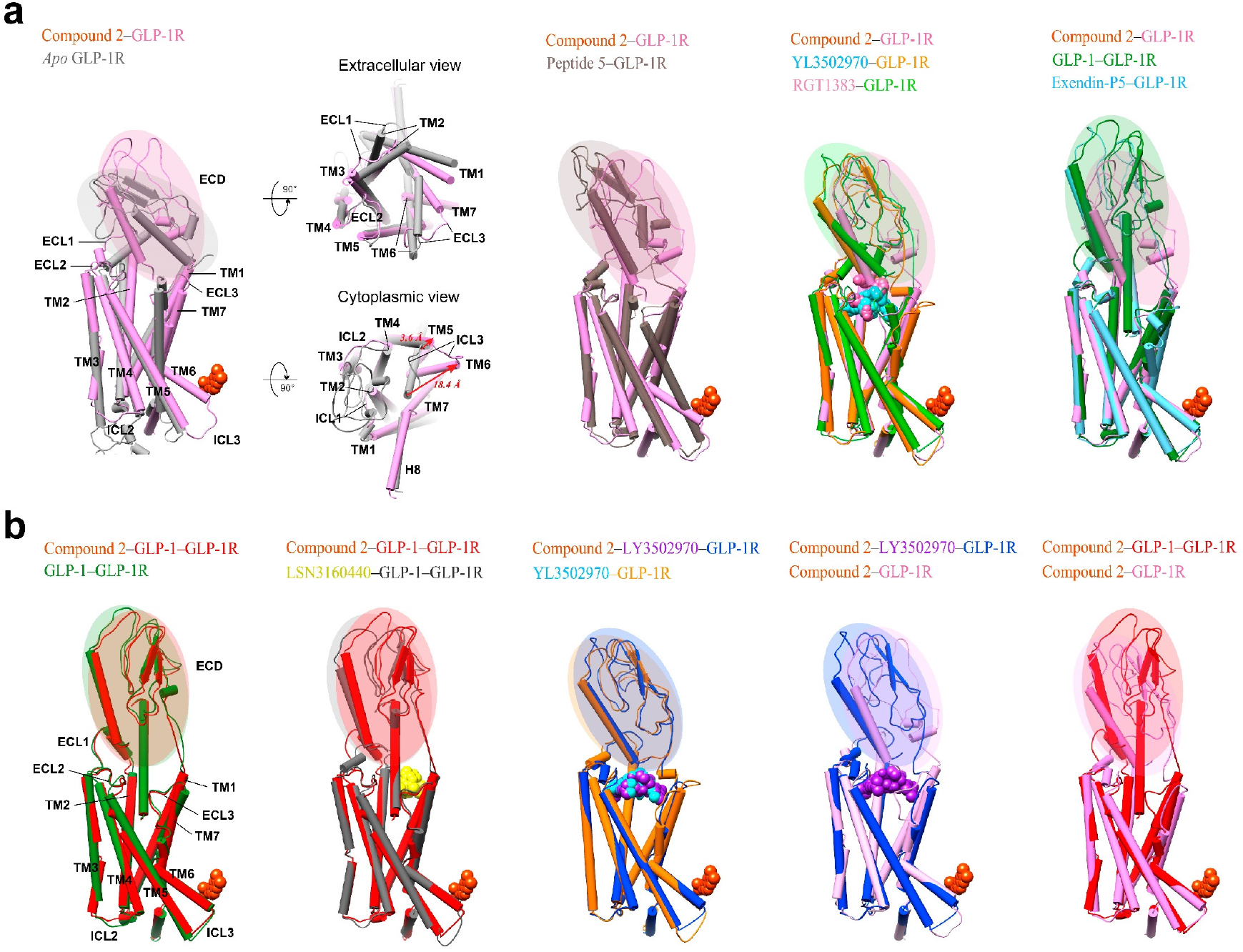
Comparison of available GLP-1R structures. **a**, Overlay of compound 2-bound GLP-1R with inactive, intermediate, peptide-bound and small molecule agonist-bound GLP-1R shows the agonism of compound 2. **b**, Overlay of compound 2 and GLP-1 or LY3502970-bound GLP-1R with related agonist bound GLP-1R shows the allosterism of compound 2. Colors of the GLP-1R and ligands are in line with the text above the structure.

**Supplementary Fig. 5.**
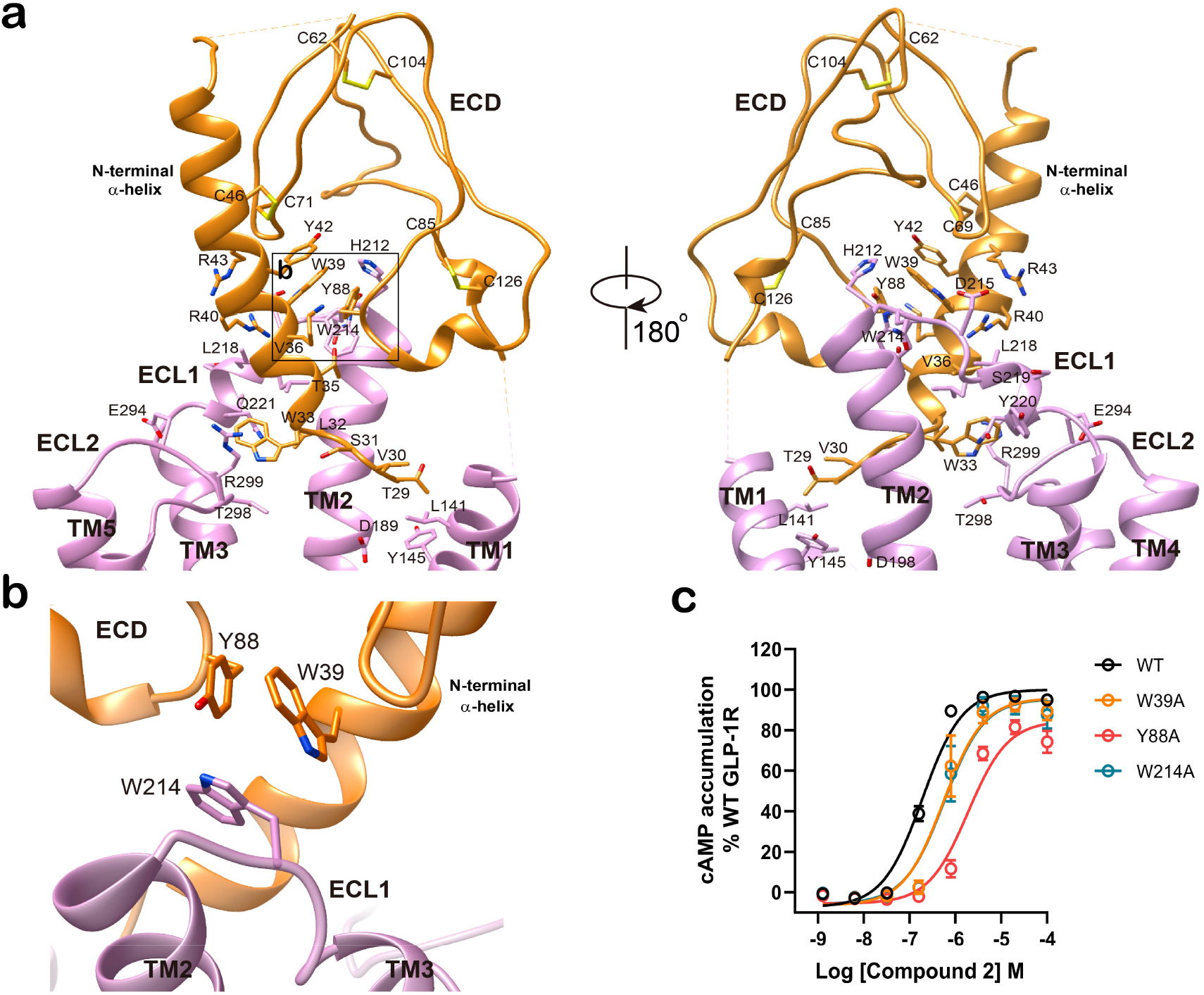
Unique ECD conformation in the compound 2-bound GLP-1R. **a**, The TMD-interacting conformation of the ECD. The ECD of GLP-1R (orange) folded down towards the TMD core and penetrated into the orthosteric binding pocket through its N-terminal α-helix. **b**, The ECD-ECL1 interactions. The ECD orientation is stabilized by interactions with ECL1. Important residues are shown in sticks. **c**, Effects of W39A, Y88A, and W214A mutants on compound 2-induced cAMP responses. Data shown are means ± S.E.M. of three independent experiments. WT, wild-type. Source data are provided as a Source Data file.

**Supplementary Fig. 6.**
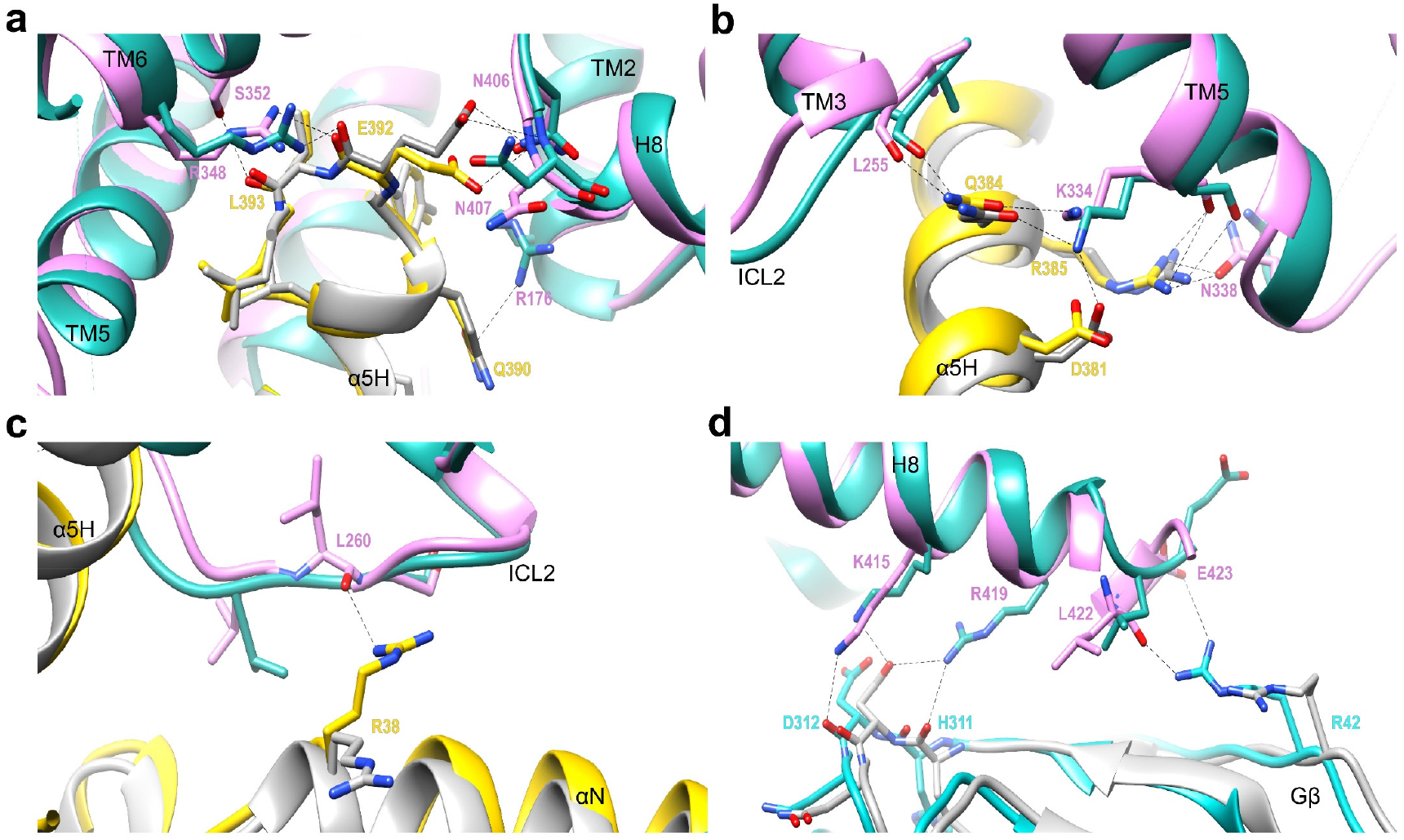
Comparison of G protein coupling between compound 2-bound and GLP-1-bound active GLP-1R in complex with G_s_. **a-b**, GLP-1R–Gα_s_ α5 helix (α5H) interface. **c**, GLP-1R–Gα_s_ N-terminal helix (αN) interface. **D**, GLP-1R helix 8-Gβ interface. Compound 2-bound GLP-1R in hot pink; GLP-1-bound GLP-1R in sea green; Gαs Ras-like domain of compound 2–GLP-1R–G_s_ complex in yellow; Gβ subunit of compound 2–GLP-1R–G_s_ complex in cyan; Gαs Ras-like domain and Gβ subunit of GLP-1–GLP-1R–G_s_ complex (PDB: 6X18) in gray.

**Supplementary Fig. 7.**
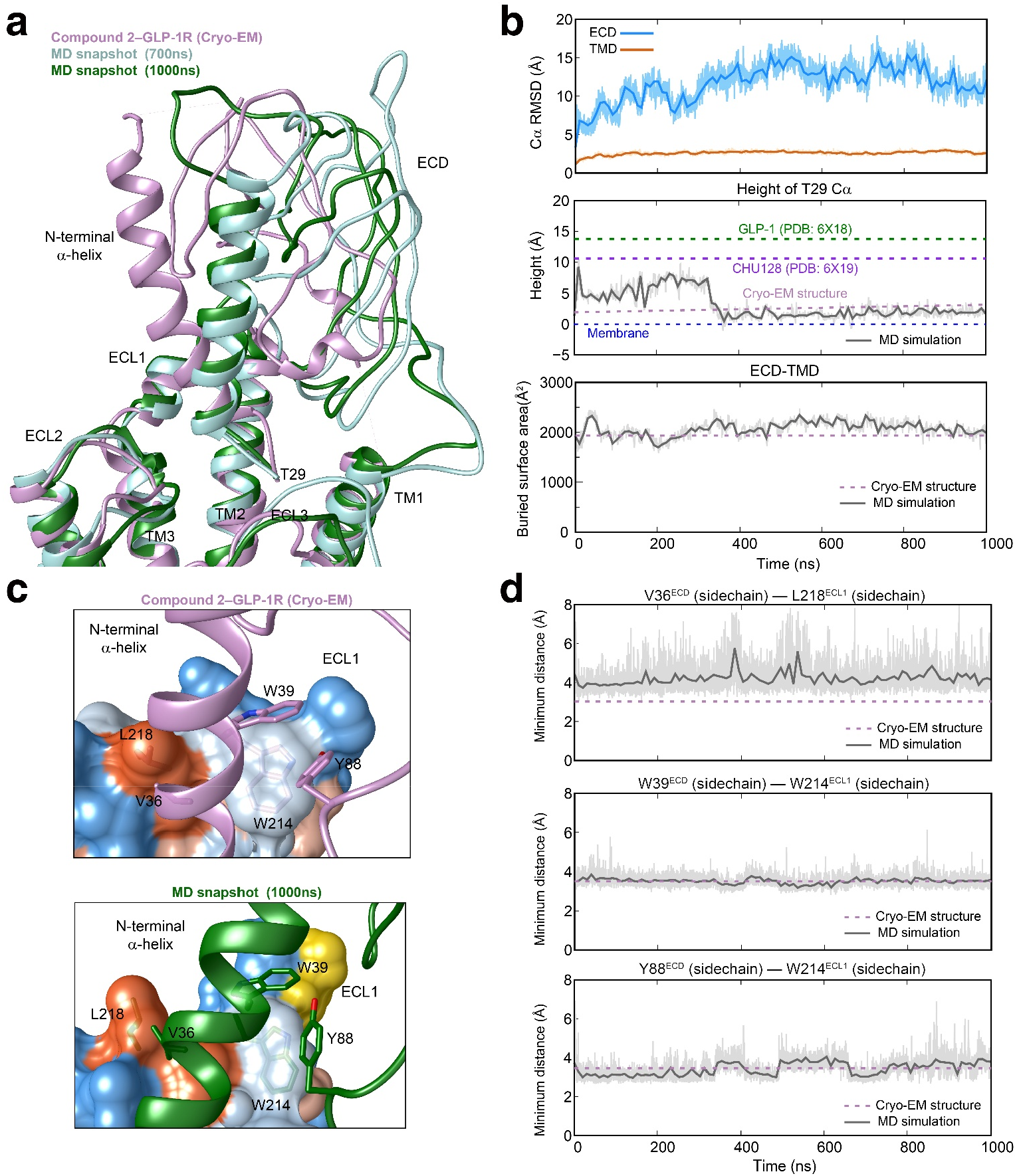
Molecular dynamics (MD) simulations of compound 2-bound active GLP-1R. **a**, Comparison of the ECD conformations between simulation snapshots and the cryo-EM structure (hot pink). **b**, Movements of the ECD during MD simulations: top, root mean square deviation (RMSD) of Cα positions of the GLP-1R, where all MD snapshots were superimposed on the cryo-EM structure of GLP-1R TMD using the Cα atoms; middle, the height of T29 Cα atom of the GLP-1R ECD relative to the membrane layer; bottom, the buried surface area between GLP-1R ECD and TMD. Interface areas were calculated using freeSASA. During the MD simulation, the N-terminal α-helix of the GLP-1R ECD consistently inserted to the TMD core, in line with the cryo-EM structure, evidenced by the height of its tip (T29) and the ECD-TMD buried surface area. The thick and thin traces represent moving averages and original, unsmoothed values, respectively. **c**, Contacts between ECD and ECL1 for the cryo-EM structure and MD snapshot. The ECL1 is shown in surface representation and colored in dodger blue for the most hydrophilic region and orange red for the most hydrophobic region, respectively. **d**, Three distances between side chain heavy atoms from the residues on the ECD and ECL1 (Top, V36^ECD^–L218^ECL1^; middle, W39^ECD^–W214^ECL1^; bottom, Y88^ECD^–W214^ECL1^). The thick and thin traces represent moving averages and original, unsmoothed values, respectively.

**Supplementary Fig. 8.**
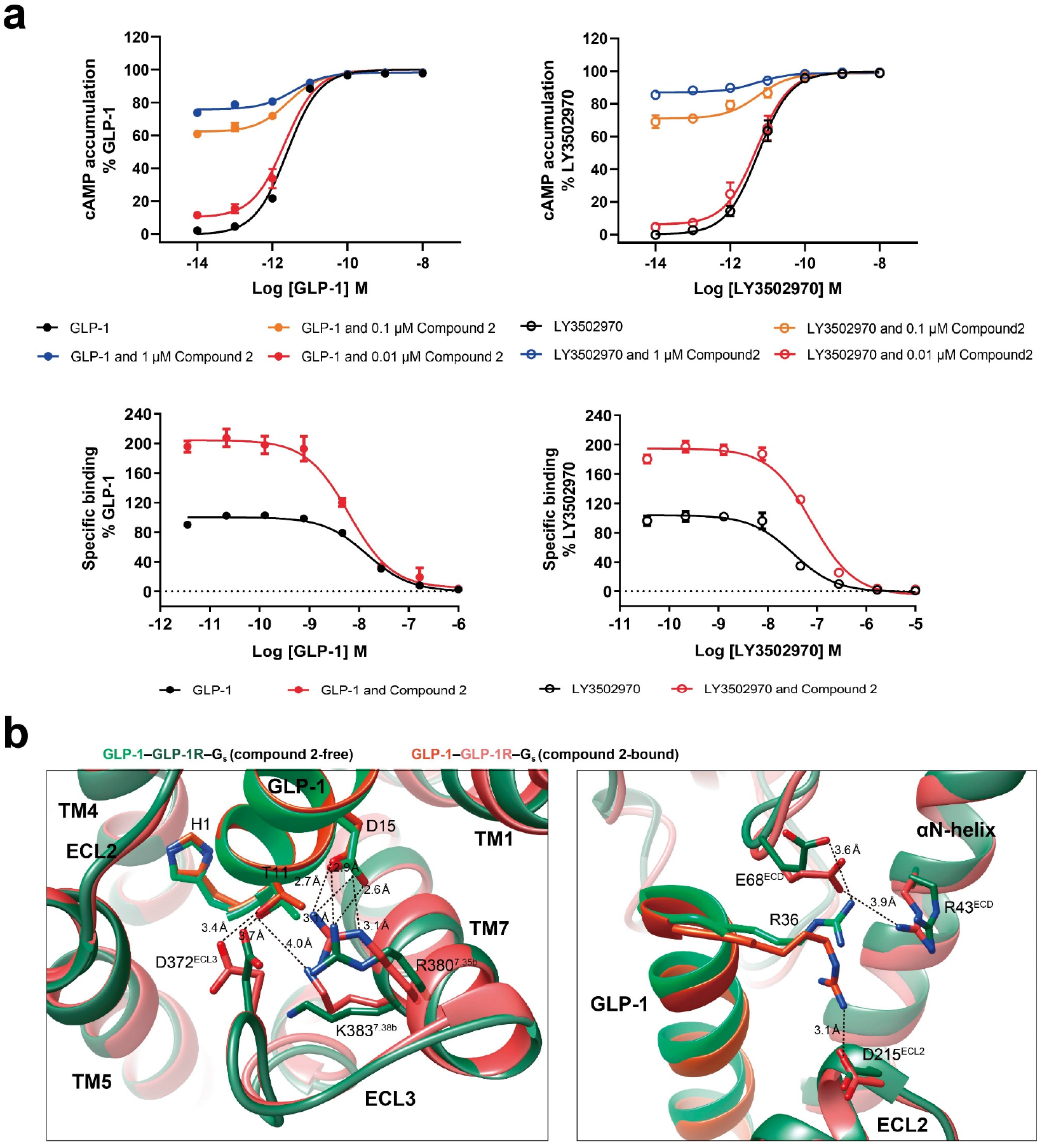
Potentiation of GLP-1 and LY3502970 activity by compound 2. **a**, Effects of compound 2 on cAMP signaling and GLP-1R binding: top, dose–response characteristics of GLP-1 and LY3502970 in the absence or presence of compound 2 at three different concentrations. In line with previous findings^24^, we did not find any increase of GLP-1 potency following of compound 2 treatment (EC50 was 2.48, 3.63, 2.63 and 2.13 pM for GLP-1 in the presence of 0, 0.1, 0.01, and 1 μM compound 2, respectively), suggesting that compound 2 only potentiates GLP-1-induced receptor activation. The same phenomenon was observed for LY3502970 (EC_50_ was 5.67, 5.31, 5.51 and 5.17 pM for LY3502970 in the presence of 0, 0.1, 0.01, and 1 μM compound 2, respectively); bottom, competitive binding of ^125^I-labelled GLP-1 with GLP-1 or LY3502970 at GLP-1R in the presence of compound 2. The results show that compound 2 enhances GLP-1 and LY3502970 binding with GLP-1R by 2-fold and 1.9-fold, respectively. Data shown are means ± S.E.M. from at least three independent experiments (n=4-6) performed in duplicate (receptor binding assay) or quadruplicate (cAMP accumulation). Source data are provided as a Source Data file. **b**, Comparison of GLP-1 and GLP-1R interaction in the presence and absence of compound 2. The cryo-EM structure of GLP-1–GLP-1R–G_s_ (PDB code: 6X18) was superimposed on Cα atoms of the compound 2–GLP-1–GLP-1R–G_s_ complex.

**Supplementary Fig. 9.**
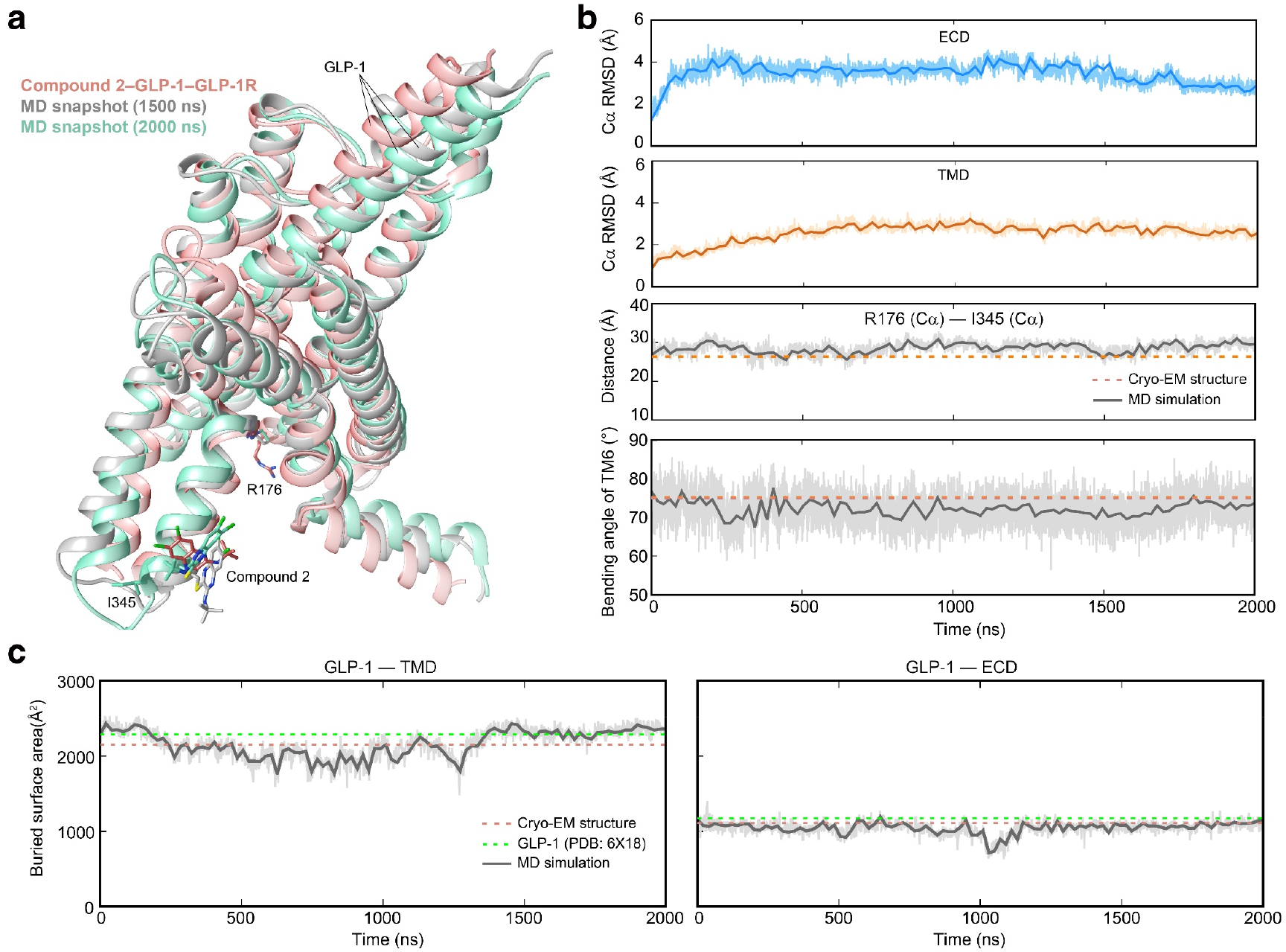
MD simulations of compound 2–GLP-1–GLP-1R. **a**, Compassion of receptor conformations between simulation snapshots and the cryo-EM structure (hot pink). ECD (residues 29 to 136) and G protein are omitted for clarity. **b**, Conformational movements of GLP-1R simulations: top, root mean square deviation (RMSD) of Cα positions of the GLP-1R ECD, which were superimposed on the ECD of the cryo-EM structure using the Cα atoms; upper middle, RMSD of Cα positions of the GLP-1R TMD (residues 137 to 423), which were superimposed on the TMD of the cryo-EM structure using the Cα atoms; lower middle, the Cα distance between two intracellular residues (R176 and I345); bottom, bending angle of TM6 (measured by the angle from L356^6.45b^ Cα to T362^6.51b^ Cα via L359^6.48b^ Cα). The thick and thin traces represent moving averages and original, unsmoothed values, respectively. **c**, The buried surface area between GLP-1 and GLP-1R. Interface areas were calculated using freeSASA. The thick and thin traces represent moving averages and original, unsmoothed values, respectively.

**Supplementary Fig. 10.**
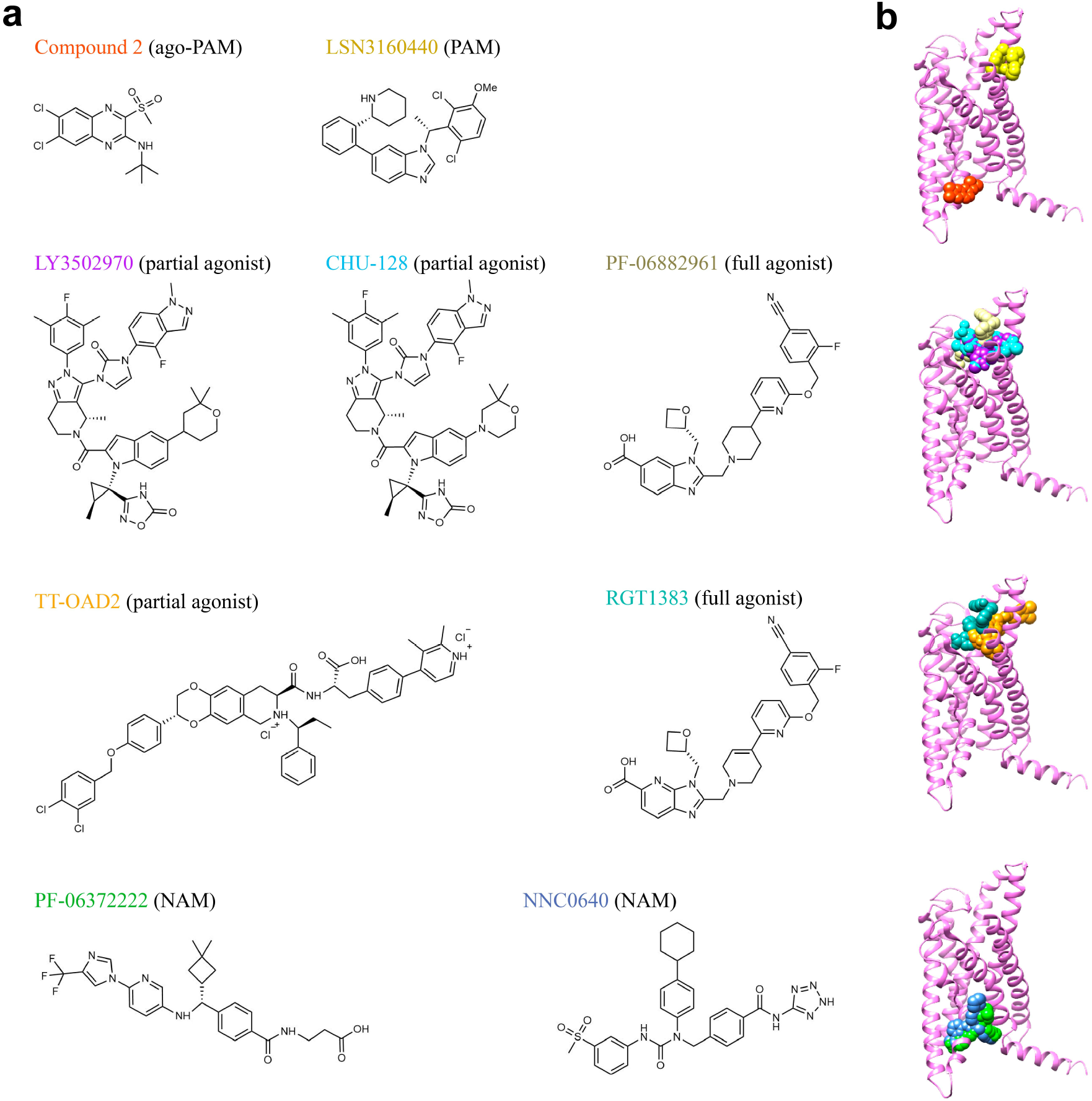
List of small molecule GLP-1R modulators with available structures. **a**, Chemical structures of small molecule ligands. **b**, Binding sites of the small molecules relative to the compound 2–GLP-1R–G_s_ complex (magenta). PAM, positive allosteric modulator; NAM, negative allosteric modulator.

**Supplementary Table. 1.** Cryo-EM data collection, refinement and validation statistics.

|  | Compound 2-GLP-1R-G<sub>s</sub><br>(without ECD) | Compound 2-GLP-1R-G<sub>s</sub><br>(with ECD) | Compound 2-GLP-1-GLP-1R-G<sub>s</sub> | Compound 2-LY3502970-GLP-1R-G<sub>s</sub> |
| --- | --- | --- | --- | --- |
| **Data collection and processing** |  |  |  |  |
| Magnification | 81000 | 81000 | 81000 | 81000 |
| Voltage (kV) | 300 | 300 | 300 | 300 |
| Electron exposure (e-/Å<sup>2</sup>) | 80 | 80 | 80 | 80 |
| Defocus range (μm) | -1.2 to -2.2 | -1.2 to -2.2 | -1.2 to -2.2 | -1.2 to -2.2 |
| Pixel size (Å) | 1.045 | 1.045 | 1.045 | 1.045 |
| Symmetry imposed | C1 | C1 | C1 | C1 |
| Initial particle images (no.) | 502,346 | 1,500,650 | 1, 664,449 | 65,661,77 |
| Final particle images (no.) | 147,173 | 340,350 | 614978 | 345411 |
| Map resolution (Å) | 3.3 | 2.5 | 2.5 | 2.9 |
| FSC threshold | 0.143 | 0.143 | 0.143 | 0.143 |
| Map resolution range (Å) | 2.8-5.0 | 2.3-5.0 | 2.2-5.0 | 2.7-5.0 |
| Map sharpening *B* factor (Å<sup>2</sup>) | -72 | -77 | -77 | -98 |
| **Refinement** |  |  |  |  |
| Initial model used (PDB code) | 6VCB | 6VCB | 6X18 | 6B3J |
| Model resolution (Å) | 2.9 | 2.9 | 2.8 | 2.9 |
| FSC threshold | 0.5 | 0.5 | 0.5 | 0.5 |
| Model resolution range (Å) | 2.3-5.0 | 2.3-5.0 | 2.2-5.0 | 2.7-5.0 |
| Model composition |  |  |  |  |
| Non-hydrogen atoms | 8,583 | 9,229 | 9,612 | 8,802 |
| Protein residues | 1,040 | 1,128 | 1,174 | 1,089 |
| Water | 0 | 0 | 0 | 0 |
| *B* factors (Å<sup>2</sup>) |  |  |  |  |
| Protein | 99.07 | 133.33 | 31.03 | 38.41 |
| Ligand | 128.62 | 99.13 | 39.12 | 36.76 |
| R.m.s. deviations |  |  |  |  |
| Bond lengths (Å) | 0.008 | 0.006 | 0.006 | 0.002 |
| Bond angles (°) | 0.687 | 0.941 | 0.729 | 0.550 |
| Validation |  |  |  |  |
| MolProbity score | 1.75 | 2.44 | 1.62 | 1.47 |
| Clashscore | 8.27 | 13.44 | 10.68 | 7.26 |
| Poor rotamers (%) | 0.22 | 0.19 | 0.20 | 0.11 |
| Ramachandran plot |  |  |  |  |
| Favored (%) | 95.61 | 95.94 | 97.66 | 97.65 |
| Allowed (%) | 4.30 | 4.81 | 2.34 | 2.35 |
| Disallowed (%) | 0.00 | 0.09 | 0.00 | 0.00 |
| Real space correlation coefficient |  |  |  |  |
| Compound 2 | 0.59 | 0.54 | 0.47 | 0.46 |
| LY3502970 |  |  |  | 0.74 |

**Supplementary Table. 2.** *In vitro* pharmacology of GLP-1R.

|  | cAMP accumulation | <sup>[125I]</sup> GLP-1 binding |  |
| --- | --- | --- | --- |
|  | GLP-1 |  |  |
|  | pEC<sub>50</sub> | E<sub>max</sub> | pKi |
| WT GLP-1R | 11.19±0.07 | 98.6±1.96 | 8.51±0.07 |
| GLP-1R-LgBiT-2MBP | 11.24±0.09 | 97.64±2.53 | 8.71±0.24 |
| Δ28 | 9.53±0.15** | 102.8±7.30 | NB** |
| Δ33 | 10.17±0.139** | 98.2±5.47 | NB** |
| Δ38 | NA** | NA** | NB** |
| Δ43 | NA** | NA** | NB** |
| Δ48 | NA** | NA** | NB** |
| Δ55 | NA** | NA** | NB** |
| ΔECD | 8.89±0.66** | 6.88±2.85** | NB** |
| V332A | 10.85±0.23 | 109.4±2.53* | 8.87±0.13* |
| C347A | 10.58±0.17 | 102.1±5.53 | 8.81±0.11 |
| K346A | 10.88±0.4 | 100.2±3.36 | 8.89±0.12* |
| L349A | 10.53±0.26* | 107.1±5.01 | 9.41±0.11** |
| A350W | 9.55±0.1** | 99.17±3.31 | 9.40±0.12** |
| K351A | 10.06±0.12** | 108.6±4.61* | 8.40±0.24 |
|  | Compound 2<sup>a</sup> |  |  |
| WT GLP-1R | 6.13±0.06 | 99.7±2.27 |  |
| GLP-1R-LgBiT-2MBP | 5.99±0.08 | 98.4±2.97 |  |
| Δ28 | 6.19±0.17 | 67.94±3.78** |  |
| Δ33 | 6.03±0.16 | 80.04±4.81** |  |
| Δ38 | 6.20±0.15 | 61.38±3.00** |  |
| Δ43 | 6.12±0.19 | 70.27±4.57** |  |
| Δ48 | 6.24±0.28 | 62.19±5.49** |  |
| Δ55 | 7.01±2.33 | 10.79±0.60** |  |
| ΔECD | 6.11±0.82 | 2.22±2.73** |  |
| V332A | 6.58±0.31 | 104.7±2.53 |  |
| C347A | 6.0±0.6 | 7.67±0.84** |  |
| K346A | 6.55±0.24 | 102.5±1.73 |  |
| L349A | 6.37±0.22 | 101.4±3.14 |  |
| A350W | 5.47±0.96 | 29.28±6.6** |  |
| K351A | 5.83±2.15 | 20.11±6.27** |  |
|  | LY3502970 |  |  |
| WT GLP-1R | 10.65±0.05 | 99.89±1.72 | 7.77±0.10 |
| GLP-1R-LgBiT-2MBP | 11.09±0.06 | 98.07±1.94 | 7.80±0.24 |

Binding data were analyzed using a three-parameter logistic equation and normalized to the maximal binding of $[^{125}\text{I}]$ GLP-1. pKi is the negative logarithm of peptide affinity. cAMP accumulation data were analyzed using a three-parameter logistic equation to determine pEC<sub>50</sub> and E<sub>max</sub> values. pEC<sub>50</sub> is the negative logarithm of the molar concentration of agonist that induced half the maximal response. E<sub>max</sub> for mutants is expressed as a percentage of the wild-type (WT). All values are means±S.E.M of at least three independent experiments conducted in duplicate. One-
way ANOVA was used to determine statistical significance (\*\*P<0.01, \*P<0.05). NA, not active; insufficient radiolabeled ligand displacement was detected implying GLP-1 binding impairment; Δ, residue truncation. Source data are provided as a Source Data file. <sup>a</sup>According to previously report<sup>24</sup>, compound 2 does not compete radiolabeled GLP-1 binding to GLP-1R and hence, it was not assessed for receptor binding.

**Supplementary Table. 3.** *In vitro* pharmacology of GCGR.

| cAMP accumulation |  |  |
| --- | --- | --- |
| Glucagon |  |  |
|  | pEC<sub>50</sub> | E<sub>max</sub> |
| WT GCGR | 8.48±0.08 | 99.39±3.68 |
| F345C | 8.43±0.08 | 95.36±3.70 |
| F345C Δ30 | 9.06±0.89** | 7.93±1.69** |
| F345C Δ35 | 8.14±0.57 | 6.9±3.27** |
| F345C Δ40 | 8.91±0.39* | 6.66±1.11** |
| F345C Δ45 | 8.28±0.45 | 4.35±1.88** |
| F345C Δ50 | 9.05±0.65 | 3.15±1.53** |
| F345C Δ55 | 8.70±0.89 | -1.93±2.79** |
| Compound 2 |  |  |
| WT GCGR | 6.74±0.83 | 11.89±1.01** |
| F345C | 6.04±0.08 | 99.87±3.16 |
| F345C Δ30 | 5.80±0.15 | 64.39±3.92** |
| F345C Δ35 | 5.88±0.20 | 52.0±4.0** |
| F345C Δ40 | 6.09±0.49 | 26.97±3.47** |
| F345C Δ45 | 5.86±0.44 | 28.43±3.73** |
| F345C Δ50 | 6.37±0.67 | 14.24±2.28** |
| F345C Δ55 | 6.03±0.45 | 24.37±3.48** |

cAMP accumulation data were analyzed using a three-parameter logistic equation to determine pEC<sub>50</sub> and E<sub>max</sub> values. pEC<sub>50</sub> is the negative logarithm of the molar concentration of agonist that induced half the maximal response. E<sub>max</sub> for mutants is expressed as a percentage of the wild-type (WT). All values are means±S.E.M of at least three independent experiments conducted in duplicate. One-way ANOVA was used to determine statistical significance (\*\*P<0.01, \*P<0.05). Δ, residue truncation. Source data are provided as a Source Data file.

**Supplementary Table. 4.**
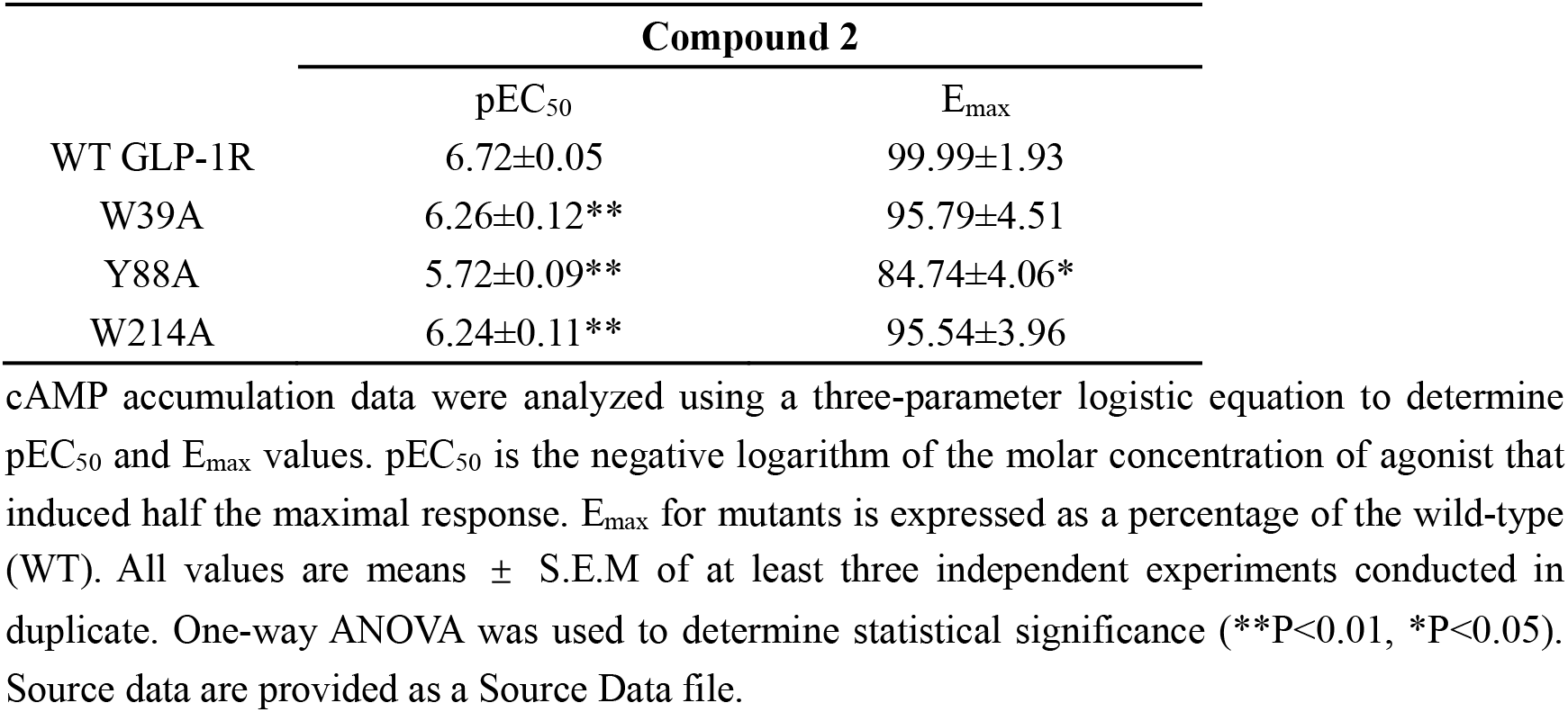
Effects of residue mutation in the ECD-ECL1 interface on cAMP accumulation.

**Supplementary Table. 5.**
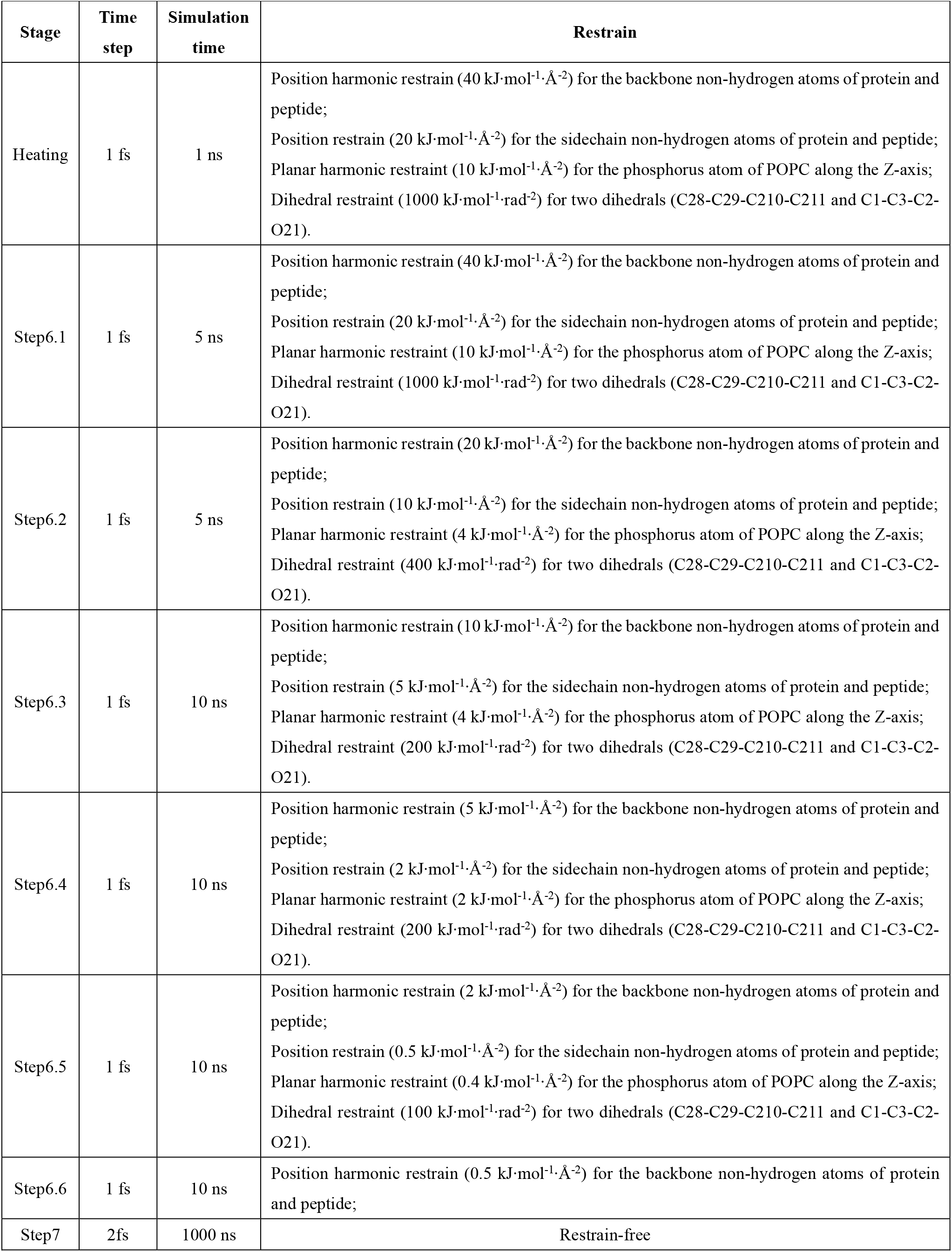
Details of restraints applied during MD simulations.

| Stage | Time step | Simulation time | Restrain |
| --- | --- | --- | --- |
| Heating | 1 fs | 1 ns | Position harmonic restrain ($40 \text{ kJ}\cdot\text{mol}^{-1}\cdot\text{\AA}^{-2}$) for the backbone non-hydrogen atoms of protein and peptide;<br>Position restrain ($20 \text{ kJ}\cdot\text{mol}^{-1}\cdot\text{\AA}^{-2}$) for the sidechain non-hydrogen atoms of protein and peptide;<br>Planar harmonic restraint ($10 \text{ kJ}\cdot\text{mol}^{-1}\cdot\text{\AA}^{-2}$) for the phosphorus atom of POPC along the Z-axis;<br>Dihedral restraint ($1000 \text{ kJ}\cdot\text{mol}^{-1}\cdot\text{rad}^{-2}$) for two dihedrals (C28-C29-C210-C211 and C1-C3-C2-O21). |
| Step6.1 | 1 fs | 5 ns | Position harmonic restrain ($40 \text{ kJ}\cdot\text{mol}^{-1}\cdot\text{\AA}^{-2}$) for the backbone non-hydrogen atoms of protein and peptide;<br>Position restrain ($20 \text{ kJ}\cdot\text{mol}^{-1}\cdot\text{\AA}^{-2}$) for the sidechain non-hydrogen atoms of protein and peptide;<br>Planar harmonic restraint ($10 \text{ kJ}\cdot\text{mol}^{-1}\cdot\text{\AA}^{-2}$) for the phosphorus atom of POPC along the Z-axis;<br>Dihedral restraint ($1000 \text{ kJ}\cdot\text{mol}^{-1}\cdot\text{rad}^{-2}$) for two dihedrals (C28-C29-C210-C211 and C1-C3-C2-O21). |
| Step6.2 | 1 fs | 5 ns | Position harmonic restrain ($20 \text{ kJ}\cdot\text{mol}^{-1}\cdot\text{\AA}^{-2}$) for the backbone non-hydrogen atoms of protein and peptide;<br>Position restrain ($10 \text{ kJ}\cdot\text{mol}^{-1}\cdot\text{\AA}^{-2}$) for the sidechain non-hydrogen atoms of protein and peptide;<br>Planar harmonic restraint ($4 \text{ kJ}\cdot\text{mol}^{-1}\cdot\text{\AA}^{-2}$) for the phosphorus atom of POPC along the Z-axis;<br>Dihedral restraint ($400 \text{ kJ}\cdot\text{mol}^{-1}\cdot\text{rad}^{-2}$) for two dihedrals (C28-C29-C210-C211 and C1-C3-C2-O21). |
| Step6.3 | 1 fs | 10 ns | Position harmonic restrain ($10 \text{ kJ}\cdot\text{mol}^{-1}\cdot\text{\AA}^{-2}$) for the backbone non-hydrogen atoms of protein and peptide;<br>Position restrain ($5 \text{ kJ}\cdot\text{mol}^{-1}\cdot\text{\AA}^{-2}$) for the sidechain non-hydrogen atoms of protein and peptide;<br>Planar harmonic restraint ($4 \text{ kJ}\cdot\text{mol}^{-1}\cdot\text{\AA}^{-2}$) for the phosphorus atom of POPC along the Z-axis;<br>Dihedral restraint ($200 \text{ kJ}\cdot\text{mol}^{-1}\cdot\text{rad}^{-2}$) for two dihedrals (C28-C29-C210-C211 and C1-C3-C2-O21). |
| Step6.4 | 1 fs | 10 ns | Position harmonic restrain ($5 \text{ kJ}\cdot\text{mol}^{-1}\cdot\text{\AA}^{-2}$) for the backbone non-hydrogen atoms of protein and peptide;<br>Position restrain ($2 \text{ kJ}\cdot\text{mol}^{-1}\cdot\text{\AA}^{-2}$) for the sidechain non-hydrogen atoms of protein and peptide;<br>Planar harmonic restraint ($2 \text{ kJ}\cdot\text{mol}^{-1}\cdot\text{\AA}^{-2}$) for the phosphorus atom of POPC along the Z-axis;<br>Dihedral restraint ($200 \text{ kJ}\cdot\text{mol}^{-1}\cdot\text{rad}^{-2}$) for two dihedrals (C28-C29-C210-C211 and C1-C3-C2-O21). |
| Step6.5 | 1 fs | 10 ns | Position harmonic restrain ($2 \text{ kJ}\cdot\text{mol}^{-1}\cdot\text{\AA}^{-2}$) for the backbone non-hydrogen atoms of protein and peptide;<br>Position restrain ($0.5 \text{ kJ}\cdot\text{mol}^{-1}\cdot\text{\AA}^{-2}$) for the sidechain non-hydrogen atoms of protein and peptide;<br>Planar harmonic restraint ($0.4 \text{ kJ}\cdot\text{mol}^{-1}\cdot\text{\AA}^{-2}$) for the phosphorus atom of POPC along the Z-axis;<br>Dihedral restraint ($100 \text{ kJ}\cdot\text{mol}^{-1}\cdot\text{rad}^{-2}$) for two dihedrals (C28-C29-C210-C211 and C1-C3-C2-O21). |
| Step6.6 | 1 fs | 10 ns | Position harmonic restrain ($0.5 \text{ kJ}\cdot\text{mol}^{-1}\cdot\text{\AA}^{-2}$) for the backbone non-hydrogen atoms of protein and peptide; |
| Step7 | 2fs | 1000 ns | Restrain-free |

**Supplementary Table. 6.**
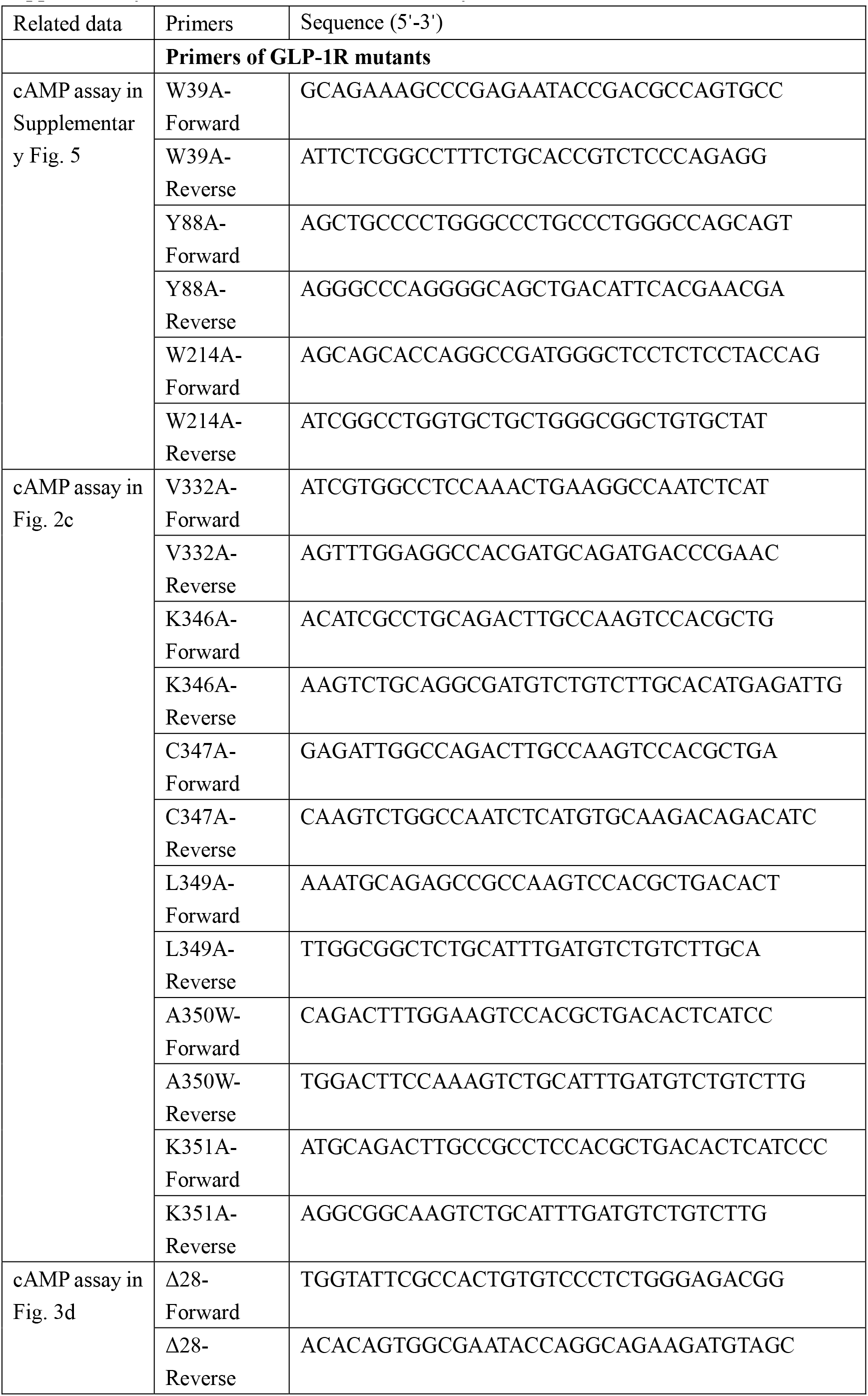
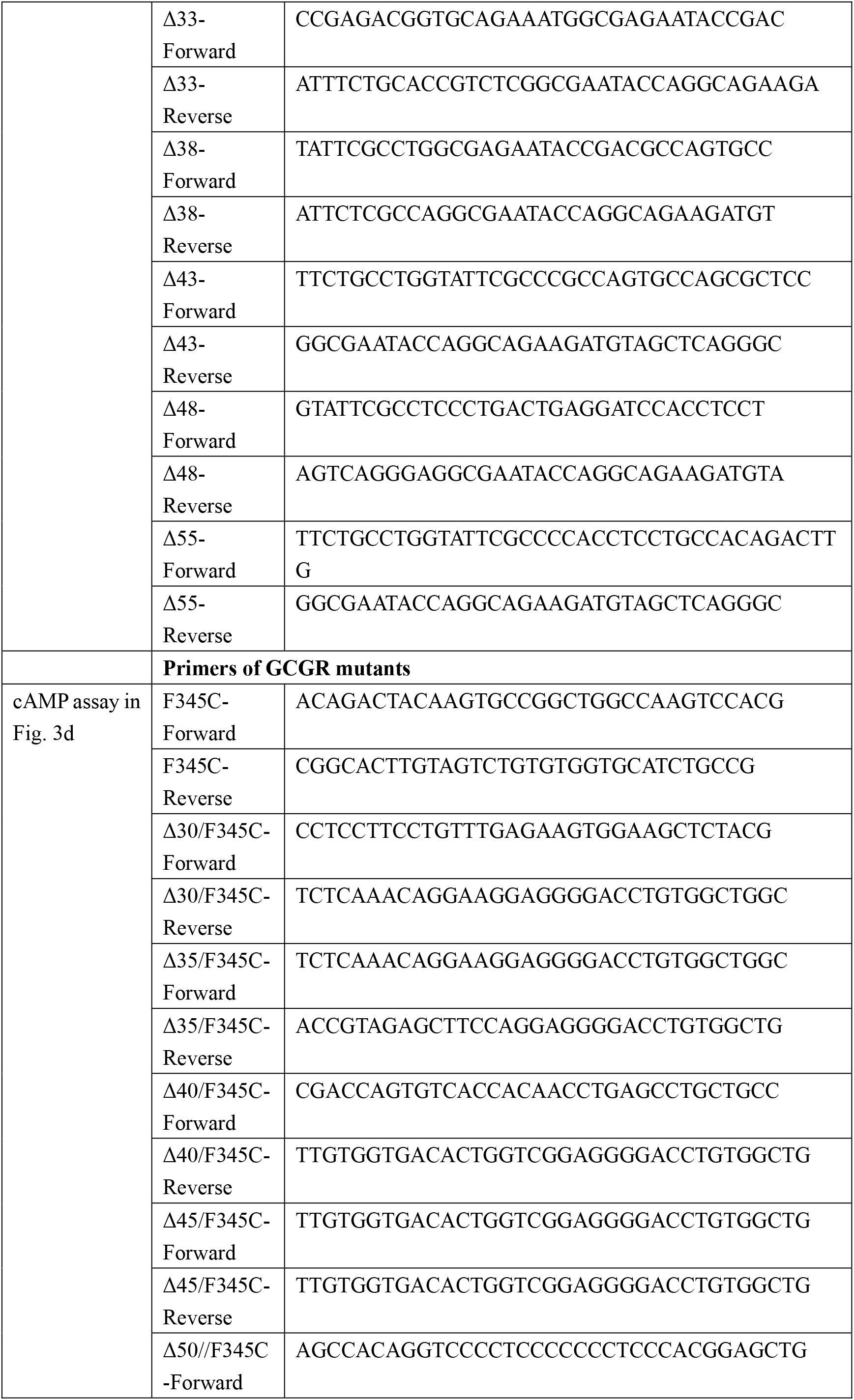
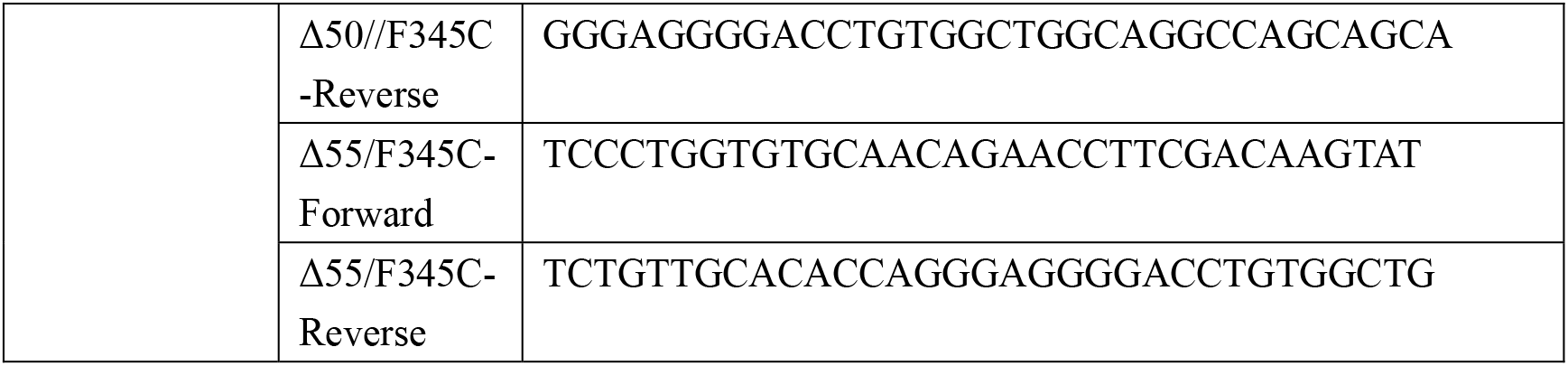
Primers used in this study.

## References

1 Saraiva, F. K. & Sposito, A. C. Cardiovascular effects of glucagon-like peptide 1 (GLP-1) receptor agonists. Cardiovasc Diabetol 13, 142, doi:10.1186/s12933-014-0142-7 (2014).

2 Drucker, D. J. Mechanisms of action and therapeutic application of glucagon-like peptide-1. Cell Metab 27, 740–756, doi:10.1016/j.cmet.2018.03.001 (2018).

3 Cho, Y. M.’ Merchant, C. E. & Kieffer, T. J. Targeting the glucagon receptor family for diabetes and obesity therapy. Pharmacol Ther 135, 247–278, doi:10.1016/j.pharmthera.2012.05.009 (2012).

4 Graaf, C. et al. Glucagon-like peptide-1 and its class B G protein-coupled receptors: a long march to therapeutic successes. Pharmacol Rev 68, 954–1013, doi:10.1124/pr.115.011395 (2016).

5 Htike, Z. Z. et al. Efficacy and safety of glucagon-like peptide-1 receptor agonists in type 2 diabetes: A systematic review and mixed-treatment comparison analysis. Diabetes Obes Metab 19’524–536, doi:10.1111/dom.12849 (2017).

6 Thethi, T. K., Pratley, R. & Meier, J. J. Efficacy, safety and cardiovascular outcomes of once-daily oral semaglutide in patients with type 2 diabetes: The PIONEER programme. Diabetes Obes Metab 22, 1263–1277, doi:10.1111/dom.14054 (2020).

7 Wright, E. E.’ Jr. & Aroda, V. R. Clinical review of the efficacy and safety of oral semaglutide in patients with type 2 diabetes considered for injectable GLP-1 receptor agonist therapy or currently on insulin therapy. Postgrad Med, 1–11, doi:10.1080/00325481.2020.1798127 (2020).

8 Liu’ C.’ Zou, Y. & Qian, H. GLP-1R agonists for the treatment of obesity: a patent review (2015-present). Expert Opin Ther Pat 30, 781–794, doi:10.1080/13543776.2020.1811851 (2020).

9 Song, G. et al. Human GLP-1 receptor transmembrane domain structure in complex with allosteric modulators. Nature 546, 312–315, doi:10.1038/nature22378 (2017).

10 Jazayeri, A. et al. Crystal structure of the GLP-1 receptor bound to a peptide agonist. Nature 546, 254–258, doi:10.1038/nature22800 (2017).

11 Zhang, Y. et al. Cryo-EM structure of the activated GLP-1 receptor in complex with a G protein. Nature 546, 248–253, doi:10.1038/nature22394 (2017).

12 Liang, Y. L. et al. Phase-plate cryo-EM structure of a biased agonist-bound human GLP-1 receptor-G_s_ complex. Nature 555, 121–125, doi:10.1038/nature25773 (2018).

13 Zhang, X. et al. Differential GLP-1R binding and activation by peptide and non-peptide agonists. Mol Cell, doi:10.1016/j.molcel.2020.09.020 (2020).

14 Zhao, P. et al. Activation of the GLP-1 receptor by a non-peptidic agonist. Nature 577, 432–436, doi:10.1038/s41586-019-1902-z (2020).

15 Ma, H. et al. Structural insights into the activation of GLP-1R by a small molecule agonist. Cell Res, doi:10.1038/s41422-020-0384-8 (2020).

16 Kawai, T. et al. Structural basis for GLP-1 receptor activation by LY3502970, an orally active nonpeptide agonist. Proc Natl Acad Sci U S A 117, 29959–29967, doi:10.1073/pnas.2014879117 (2020).

17 Bueno, A. B. et al. Structural insights into probe-dependent positive allosterism of the GLP-1 receptor. Nat Chem Biol 16, 1105–1110, doi:10.1038/s41589-020-0589-7 (2020).

18 Wu, F. et al. Full-length human GLP-1 receptor structure without orthosteric ligands. Nat Commun 11, 1272, doi:10.1038/s41467-020-14934-5 (2020).

19 Felder, C. C. GPCR drug discovery-moving beyond the orthosteric to the allosteric domain. Adv Pharmacol 86, 1–20, doi:10.1016/bs.apha.2019.04.002 (2019).

20 Wootten, D. & Miller, L. J. Structural basis for allosteric modulation of class B G protein-coupled receptors. Annu Rev Pharmacol Toxicol 60, 89–107, doi:10.1146/annurev-pharmtox-010919-023301 (2020).

21 Malik, F. & Li, Z. Non-peptide agonists and positive allosteric modulators of glucagon-like peptide-1 receptors: Alternative approaches for treatment of Type 2 diabetes. Br J Pharmacol,doi:10.1111/bph.15446 (2021).

22 Willard, F. S. et al. Discovery of an orally efficacious positive allosteric modulator of the glucagon-like peptide-1 receptor. J Med Chem 64, 3439–3448, doi: 10.1021/acsjmedchem.1c00029 (2021).

23 Sloop, K. W. et al. Novel small molecule glucagon-like peptide-1 receptor agonist stimulates insulin secretion in rodents and from human islets. Diabetes 59, 3099–3107, doi:10.2337/db10-0689 (2010).

24 Bueno, A. B. et al. Positive allosteric modulation of the glucagon-like peptide-1 receptor by diverse electrophiles. J Biol Chem 291, 10700–10715, doi:10.1074/jbc.M115.696039 (2016).

25 Knudsen, L. B. et al. Small-molecule agonists for the glucagon-like peptide 1 receptor. Proc Natl Acad Sci U S A 104, 937–942, doi:10.1073/pnas.0605701104 (2007).

26 Koole, C. et al. Differential impact of amino acid substitutions on critical residues of the human glucagon-like peptide-1 receptor involved in peptide activity and small-molecule allostery. J Pharmacol Exp Ther 353, 52–63, doi:10.1124/jpet.114.220913 (2015).

27 Nolte, W. M. et al. A potentiator of orthosteric ligand activity at GLP-1R acts via covalent modification. Nat Chem Biol 10, 629–631, doi:10.1038/nchembio.1581 (2014).

28 Cheong, Y. H., Kim, M. K., Son, M. H. & Kaang, B. K. Two small molecule agonists of glucagon-like peptide-1 receptor modulate the receptor activation response differently. Biochem Biophys Res Commun 417, 558–563, doi:10.1016/j.bbrc.2011.12.004 (2012).

29 Wootten, D. et al. Differential activation and modulation of the glucagon-like peptide-1 receptor by small molecule ligands. Mol Pharmacol 83, 822–834, doi:10.1124/mol.112.084525 (2013).

30 Coopman, K. et al. Comparative effects of the endogenous agonist glucagon-like peptide-1 (GLP-1)-(7-36) amide and the small-molecule ago-allosteric agent “compound 2” at the GLP-1 receptor. J Pharmacol Exp Ther 334, 795–808, doi:10.1124/jpet.110.166009 (2010).

31 Koole, C. et al. Allosteric ligands of the glucagon-like peptide 1 receptor (GLP-1R) differentially modulate endogenous and exogenous peptide responses in a pathway-selective manner: implications for drug screening. Mol Pharmacol 78, 456–465, doi:10.1124/mol.110.065664 (2010).

32 Wootten, D. et al. Allosteric modulation of endogenous metabolites as an avenue for drug discovery. Mol Pharmacol 82, 281–290, doi:10.1124/mol.112.079319 (2012).

33 Willard, F. S., Ho, J. D. & Sloop, K. W. Discovery and pharmacology of the covalent GLP-1 receptor (GLP-1R) allosteric modulator BETP: A novel tool to probe GLP-1R pharmacology. Adv Pharmacol 88, 173–191, doi:10.1016/bs.apha.2020.02.001 (2020).

34 Zhao, L. H. et al. Differential Requirement of the Extracellular domain in activation of class B G protein-coupled receptors. J Biol Chem 291, 15119–15130, doi:10.1074/jbc.M116.726620 (2016).

35 Duan, J. et al. Cryo-EM structure of an activated VIP1 receptor-G protein complex revealed by a NanoBiT tethering strategy. Nat Commun 11, 4121, doi:10.1038/s41467-020-17933-8 (2020).

36 Zhou, F. et al. Structural basis for activation of the growth hormone-releasing hormone receptor. Nat Commun 11, 5205, doi:10.1038/s41467-020-18945-0 (2020).

37 Wootten, D., Simms, J., Miller, L. J., Christopoulos, A. & Sexton, P. M. Polar transmembrane interactions drive formation of ligand-specific and signal pathway-biased family B G protein-coupled receptor conformations. Proc Natl Acad Sci U S A 110, 5211–5216, doi:10.1073/pnas.1221585110 (2013).

38 Teng, M. et al. Small molecule ago-allosteric modulators of the human glucagon-like peptide-1 (hGLP-1) receptor. Bioorg Med Chem Lett 17, 5472–5478, doi:10.1016/j.bmcl.2007.06.086 (2007).

39 Willard, F. S. & Sloop, K. W. Physiology and emerging biochemistry of the glucagon-like peptide-1 receptor. Exp Diabetes Res 2012, 470851, doi:10.1155/2012/470851 (2012).

40 Yin, Y. et al. An intrinsic agonist mechanism for activation of glucagon-like peptide-1 receptor by its extracellular domain. Cell Discov 2, 16042, doi:10.1038/celldisc.2016.42 (2016).

41 Yang, L. et al. Conformational states of the full-length glucagon receptor. Nat Commun 6, 7859, doi:10.1038/ncomms8859 (2015).

42 Mattedi, G., Acosta-Gutierrez, S., Clark, T. & Gervasio, F. L. A combined activation mechanism for the glucagon receptor. Proc Natl Acad Sci U S A 117, 15414–15422, doi:10.1073/pnas.1921851117 (2020).

43 Hoare, S. R. Mechanisms of peptide and nonpeptide ligand binding to Class B G-protein-coupled receptors. Drug Discov Today 10, 417–427, doi:10.1016/S1359-6446(05)03370-2 (2005).

44 Schwartz, T. W. & Holst, B. Ago-allosteric modulation and other types of allostery in dimeric 7TM receptors. J Recept Signal Transduct Res 26, 107–128, doi:10.1080/10799890600567570 (2006).

45 Hilger, D. et al. Structural insights into differences in G protein activation by family A and family B GPCRs. Science 369, doi:10.1126/science.aba3373 (2020).

46 Yin, Y. et al. Rearrangement of a polar core provides a conserved mechanism for constitutive activation of class B G protein-coupled receptors. J Biol Chem 292, 9865–9881, doi:10.1074/jbc.M117.782987 (2017).

47 Liang, Y. L. et al. Dominant negative G proteins enhance formation and purification of agonist-GPCR-G protein complexes for structure determination. ACS Pharmacol Transl Sci 1, 12–20, doi:10.1021/acsptsci.8b00017 (2018).

48 Carpenter, B., Nehme, R., Warne, T., Leslie, A. G. & Tate, C. G. Structure of the adenosine A(2A) receptor bound to an engineered G protein. Nature 536, 104–107, doi:10.1038/nature18966 (2016).

49 Kang, Y. et al. Cryo-EM structure of human rhodopsin bound to an inhibitory G protein. Nature 558, 553–558, doi:10.1038/s41586-018-0215-y (2018).

50 Maeda, S. et al. Development of an antibody fragment that stabilizes GPCR/G-protein complexes. Nat Commun 9, 3712, doi:10.1038/s41467-018-06002-w (2018).

51 Maeda, S., Qu, Q., Robertson, M. J., Skiniotis, G. & Kobilka, B. K. Structures of the M1 and M2 muscarinic acetylcholine receptor/G-protein complexes. Science 364, 552–557, doi:10.1126/science.aaw5188 (2019).

52 Rasmussen, S. G. et al. Crystal structure of the beta2 adrenergic receptor-G_s_ protein complex. Nature 477, 549–555, doi:10.1038/nature10361 (2011).

53 Zheng, S. Q. et al. MotionCor2: anisotropic correction of beam-induced motion for improved cryo-electron microscopy. Nat Methods 14, 331–332, doi:10.1038/nmeth.4193 (2017).

54 Zhang, K. Gctf: Real-time CTF determination and correction. J Struct Biol 193, 1–12, doi:10.1016/j.jsb.2015.11.003 (2016).

55 Scheres, S. H. RELION: implementation of a Bayesian approach to cryo-EM structure determination. J Struct Biol 180, 519–530, doi:10.1016/j.jsb.2012.09.006 (2012).

56 Heymann, J. B. Guidelines for using Bsoft for high resolution reconstruction and validation of biomolecular structures from electron micrographs. Protein Sci 27, 159–171, doi:10.1002/pro.3293 (2018).

57 Wu, E. L. et al. CHARMM-GUI Membrane Builder toward realistic biological membrane simulations. J Comput Chem 35, 1997–2004, doi:10.1002/jcc.23702 (2014).

58 Guvench, O. et al. CHARMM additive all-atom force field for carbohydrate derivatives and its utility in polysaccharide and carbohydrate-protein modeling. J Chem Theory Comput 7, 3162–3180, doi:10.1021/ct200328p (2011).

59 Yu, W., He, X., Vanommeslaeghe, K. & MacKerell, A. D., Jr. Extension of the CHARMM General Force Field to sulfonyl-containing compounds and its utility in biomolecular simulations. J Comput Chem 33, 2451–2468, doi:10.1002/jcc.23067 (2012).

60 Hess, B. P-LINCS: a parallel linear constraint solver for molecular simulation. J Chem Theory Comput 4, 116–122, doi:10.1021/ct700200b (2008).

61 Aoki, K. M. & Yonezawa, F. Constant-pressure molecular-dynamics simulations of the crystalsmectic transition in systems of soft parallel spherocylinders. Phys Rev A 46, 6541–6549, doi:10.1103/physreva.46.6541 (1992).

62 Mitternacht, S. FreeSASA: An open source C library for solvent accessible surface area calculations. F1000Res 5, 189, doi:10.12688/f1000research.7931.1 (2016).

